# Energetic analysis of Na^+^/K^+^-ATPase using bond graphs

**DOI:** 10.64898/2026.04.05.716446

**Authors:** Weiwei Ai, Peter J. Hunter, Michael Pan, David P. Nickerson

## Abstract

The sodium-potassium ATPase (NKA) accounts for 19–28% of the total ATP consumption in mammals and is critical for maintaining ion homeostasis. Understanding its energetic efficiency is essential for comprehending cellular physiology and pathophysiology. We develop bond graph models of the NKA that ensure thermodynamic consistency by enforcing conservation of mass, charge, and energy. A simplified 6-state model captures biophysics comparable to a 15-state model while remaining computationally tractable. Through detailed energetic analysis, we demonstrate that under physiological conditions for the mammalian cardiac cell, approximately 66% of the energy from ATP hydrolysis is stored as chemical energy in ion gradients, 12% as electrical energy in the membrane potential, and 22% is dissipated as heat, yielding an overall efficiency of ~78%. We investigate how the free energy of ATP hydrolysis (Δ*G*_*ATP*_), intracellular Na^+^, and extracellular K^+^ affect NKA efficiency and activity. A critical threshold exists at Δ*G*_*ATP*_ ≈ −48 kJ/mol below which chemoelectrical transduction drops dramatically, consistent with NKA inhibition under ischemic conditions. The bond graph framework enables quantitative comparison of different NKA models and provides a systematic approach for analyzing ion pumps.

**SIGNIFICANCE:** The sodium-potassium ATPase is one of the body’s most energy-consuming enzymes, yet its energetic efficiency and mechanisms remain incompletely understood. This study presents the first comprehensive energetic analysis using bond graph modeling, guaranteeing thermodynamic consistency. By demonstrating that simplified 6-state models capture essential energetic behaviors of complex 15-state models, we establish bond graphs as a powerful, tractable tool for energetic analysis, model comparison, model selection and validation. The bond graph approach can be applied to other transporters, offering a powerful tool for systems physiology and drug discovery.

## INTRODUCTION

The sodium-potassium ATPase pump (NKA) is a ubiquitous enzyme that uses the energy of the hydrolysis of adenosine triphosphate (ATP) to move Na^+^ ions out of the cell and K^+^ ions into the cell, in both cases against steep concentration gradients. ATP hydrolysis can be thought of as the energy source that tops up the cell’s ‘sodium’ battery, since the majority of transmembrane transport processes are driven by this sodium gradient. The NKA is therefore critical to cellular function in all cell types and relevant to various pathophysiological conditions (1).

In this paper we use a bond graph approach to examine a number of models of NKA in which mass, charge and energy are each conserved. We examine a previously published 15-state bond graph model, and then show how a much simpler 6-state model captures the biophysics of the pump and matches experimental data. The reaction schemes of these two models are both based on the Albers-Post scheme (2, 3), but the reaction steps and orders are quite different. Usually, the comparison of different models is based on the steady state fluxes of the models, while the bond graph approach allows us to directly compare the energetic behavior of different models under dynamic conditions. We examine the energy expenditure of the NKA and energy storage and dissipation under different conditions. These energetic analyses are important to understand the efficiency of the NKA and its role in cellular energetics. We also discuss how the energetic analysis can provide insights into the physiological function of the NKA and its implications for cellular health and disease. We show how energetic profiles of NKA under physiological and pathophysiological conditions can be used for model validation, model comparison and model selection for different applications.

### NKA physiology and models

The NKA was discovered in 1957(4), and is a chemical molecular machine responsible for the movement of Na^+^ and K^+^ ions across the cell membrane against their concentration gradients. The Albers–Post kinetic scheme (2, 3) was proposed to describe the transport mechanism of the NKA (5), where the NKA undergoes Na^+^-dependent phosphorylation and K^+^-dependent dephosphorylation reactions and alternates between two main conformations, E1 and E2, which expose the transport sites to the intracellular and extracellular sides of the membrane, respectively.

The cryo-electron microscopy (cryo-EM) structures of NKA in different conformational states were reported, including exoplasmic side-open (E2·P)(6, 7), K^+^-occluded (E2·[2K] ·Pi) (7–9), (E2·[2K])(10), cytoplasmic side-open (E1)(7), ATP bound cytoplasmic side-open (E1·ATP)(7), (E1·[3Na]·ATP)(10), Na^+^-occluded (E1·[3Na]·P-ADP) (7, 11, 12), and (E1·[3Na])(10). While these structural studies have provided insights into the molecular mechanism of NKA, Wagoner et al. (13) discovered that NKA achieved high speed and efficiency by utilizing these multiple conformational states.

Since the NKA transports three Na^+^ ions outward and two K^+^ ions inward, the process gives rise to a net outward current and is therefore electrogenic (5, 14). That is, the NKA contributes to the membrane potential and the transport rate is a function of the membrane potential. The electrogenicity of the NKA has been studied experimentally and the main voltage-dependent steps are associated with the binding and release of Na^+^ ions (15–21). More recently, the electrogenicity of extracellular K^+^ binding was also investigated (22, 23) while K^+^ uptake was found to be less voltage-dependent (23, 24).

Many kinetic NKA models, including a varied number of intermediate conformational states and partial reaction steps, have been proposed (5, 25–29). Most of these models are based on the Albers-Post scheme (2, 3). Terkildsen et al. (28) developed a kinetic model of the NKA including fifteen intermediate states, also based on the Albers-Post scheme. This model detailed the binding and release of Na^+^ and K^+^ ions and their voltage dependence. It also explicitly included the proton to enable the study of the effect of pH on pump function, which was found to be significant in ischemic conditions. The model was subsequently corrected by Pan et al. (30) to address inconsistencies in the original model using a bond graph approach to ensure thermodynamic consistency.

The NKA consumes 19-28% of the total ATP consumption in mammals, and up to 50-60% in the brain and kidney (31). The experimental study in guinea-pig cardiac ventricular muscle showed that the NKA is responsible for 10% of myocardial heat production during contraction, and 17% at rest (32). It is therefore important to guarantee thermodynamic consistency in NKA model development so that heat production can be quantified and compared to available experimental data. Hence, the bond graph is also used in this study, and we demonstrate how this approach facilitates energy-based analysis.

### Bond graph in systems biology

Bond graphs are an energy-based modelling framework originally developed in the field of engineering (33–35). A comprehensive review of bond graphs in systems biology can be found in (36).

Gawthrop and Crampin (37) pioneered the application of bond graphs to biochemical systems, demonstrating how bond graphs can ensure thermodynamic consistency in biochemical network models. They showed that bond graphs can be used to model metabolic networks while adhering to the laws of thermodynamics, and to facilitate model reduction while preserving structural and thermodynamic properties. The modularity of bond graphs was further explored to enable model composition with guaranteed thermodynamic consistency in (38–41). The bond graph has been extended to include chemoelectrical transduction (42), which illustrates how the energy-based approach can explicitly account for the energy flows in excitable membranes. Pan et al. (43, 44) developed bond graph models of cardiac action potentials, demonstrating how the bond graph framework can be used to examine the conservation of physics and explain the model drift and non-unique steady states. In (45), they demonstrated how bond graphs can facilitate analysis of energy transduction in single transporters. More complex bond graph models with feedback control have been used in (46), where the authors integrate linear control theory with bond graph modelling to analyse energy flows in biochemical networks with cyclic flow modulation. These studies highlight the versatility and power of bond graphs in systems biology, particularly in ensuring thermodynamic consistency and facilitating energy-based analysis (47).

## METHODS

### The bond graph modelling framework

Bond graphs provide a useful way of formulating and visualising a thermodynamically valid biophysical model because they ensure that mass, charge and energy are each conserved, and they clearly distinguish the mechanisms for (i) transmission of power, (ii) energy storage (mechanically in a spring, electrically in a capacitor or chemically in a solute dissolved in a solvent), (iii) energy dissipation to heat (a mechanical damper, an electrical resistance or a chemical reaction), and (iv) energy transfer between mechanical, electrical or biochemical domains. Most importantly they distinguish between the conservation laws of physics and the empirically measured constitutive parameters that characterise particular materials.

In this article, we use a slightly different notation to that used in previous bond graph papers (30, 37, 43, 44, 48) to make the bond graph more accessible to a wider audience. We refer the reader to (30, 37, 43, 44, 48) for more details of the bond graph methodology. Here, we only briefly introduce the key concepts of bond graph modelling relevant to this paper and explain the new notation used in the bond graph models of NKA, as shown in Figure 1. Lines of power transmission called *bonds* always have an associated flux *v* and potential *u*, and the product of flux *v* and potential *u* is power, i.e., *P* = *u* · *v*. The arrow direction specifies the positive reference direction of the energy flow, i.e., the direction of power flow (Figure 1(a)). In a dynamic system, the energy flow *P*(*t*) = *u*(*t*) · *v*(*t*) direction may change over time, while a positive power *P*(*t*) > 0 indicates the energy is released at the tail at time *t*.

**Figure 1:**
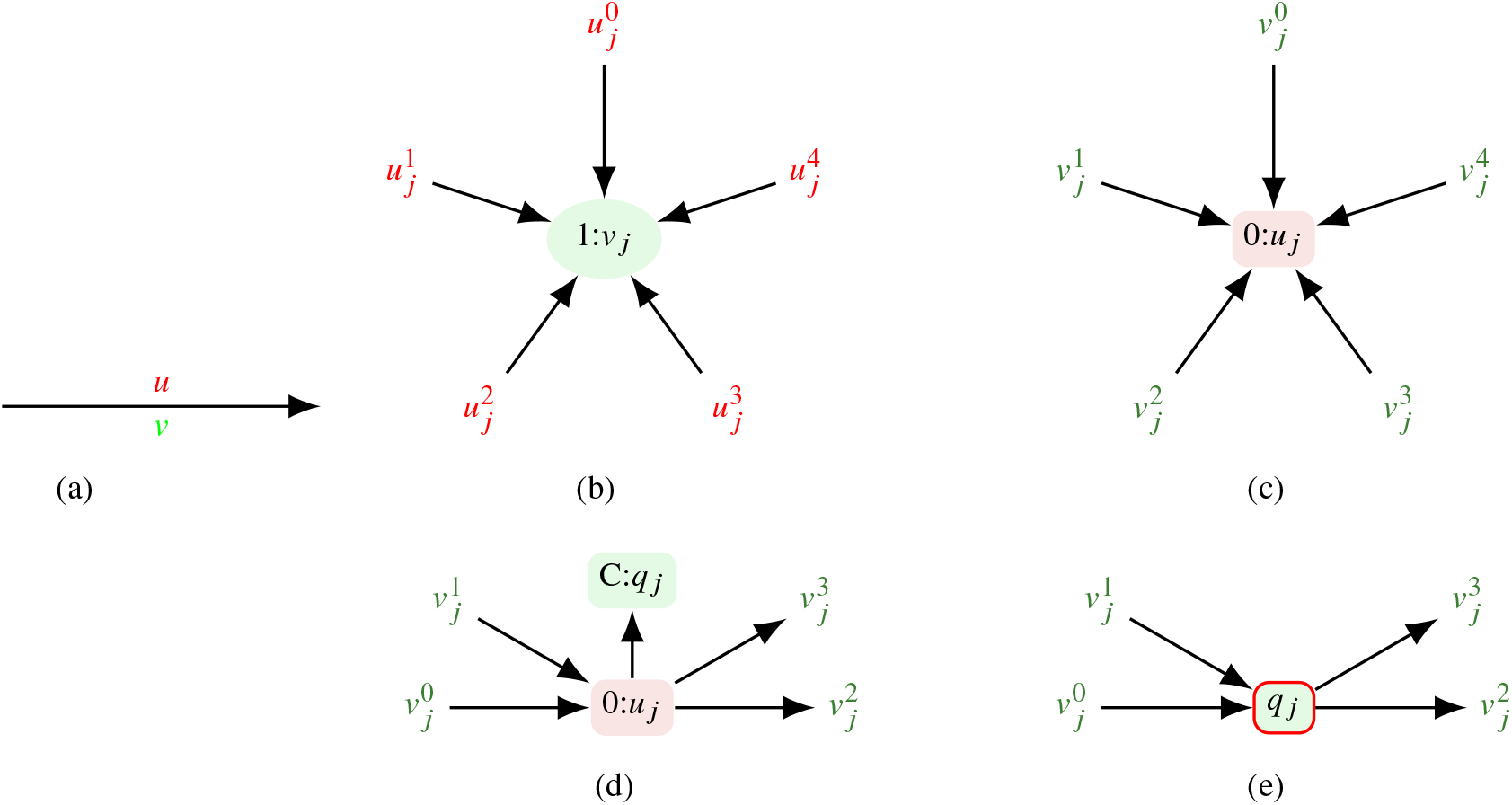
(a) a bond, which transmits energy, carries a flow *v*_*j*_ and a potential *u*_*j*_; (b) a 1 node is a bond junction, where the flow is the same for all bonds and therefore the sum of potentials is zero (conservation of energy); (c) a 0 node is a bond junction, where the potential is the same for all bonds and therefore the sum of flows is zero (conservation of mass or charge); (d) a 0 node is usually associated with capacitive energy storage; (e) a concise way to be more succinctly expressed by the red-bordered box where the potential *u*_*j*_ is given by an empirically defined capacitive storage relationship *u*_*j*_ = *f*(*q*_*j*_).

If these bonds meet, the sum of powers must be zero ∑*u* · *v* = 0, to ensure power conservation. If they share a common flux *v* (called a *1 node*), power conservation becomes just ∑*u* = 0 (for non-zero v), which is energy conservation (Figure 1(b)). Alternatively, if they share a common potential *u* (called a *0 node*), power conservation becomes just ∑*v* = 0 (for non-zero u), which is mass conservation if *v* is a molar flux or mechanical flow and charge conservation if *v* is an electrical flux (Figure 1(c)).

The explicit expression of energy flows in bond graphs makes them particularly suitable for thermodynamic analysis of biochemical systems. For each component *j* in the bond graph, the energy flow can be calculated as the sum of the power transmitted by each of its bonds, that is 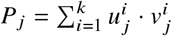 with *i* = 1, 2, …, *k* indicating the number of bonds connected to the component. For dynamic systems, the power can be integrated over time to give the total energy stored or dissipated by the component over the time interval *t*_0_ to *t*_1_, that is 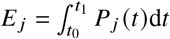.

The total energy storage and dissipation of the system can then be determined by summing the energy flows of all components. This allows for a detailed analysis of the energy budget of the system, including identifying which components are storing or dissipating the most energy, and how changes in system parameters affect the overall energy balance.

### Bond graph models of NKA

Here, we explain the construction of a 6-state bond graph model of NKA using the new notation, while the 15-state model is explained in (30) and the graph using the new notation is in the supplementary material. The bond graph model of NKA with 6 states is based on the scheme proposed in Fig. 1b of (7) with slight modifications. We merged (E1·[3Na]·ATP) and (E1·[3Na]·P-ADP), while splitting the release of adenosine diphosphate (ADP) and Na^+^ into two steps. The graphic representation of the model is shown in Figure 2 with the steps of the transporters, where we label the cytoplasmic side-open state (E1) as *E*_*i*_ and the exoplasmic side-open state E2 as *E*_*o*_. We also derived the analytical steady state equations of this model in the supplementary material, which will be used to check when the instantaneous dynamics of the model differ from the steady state.

**Figure 2:**
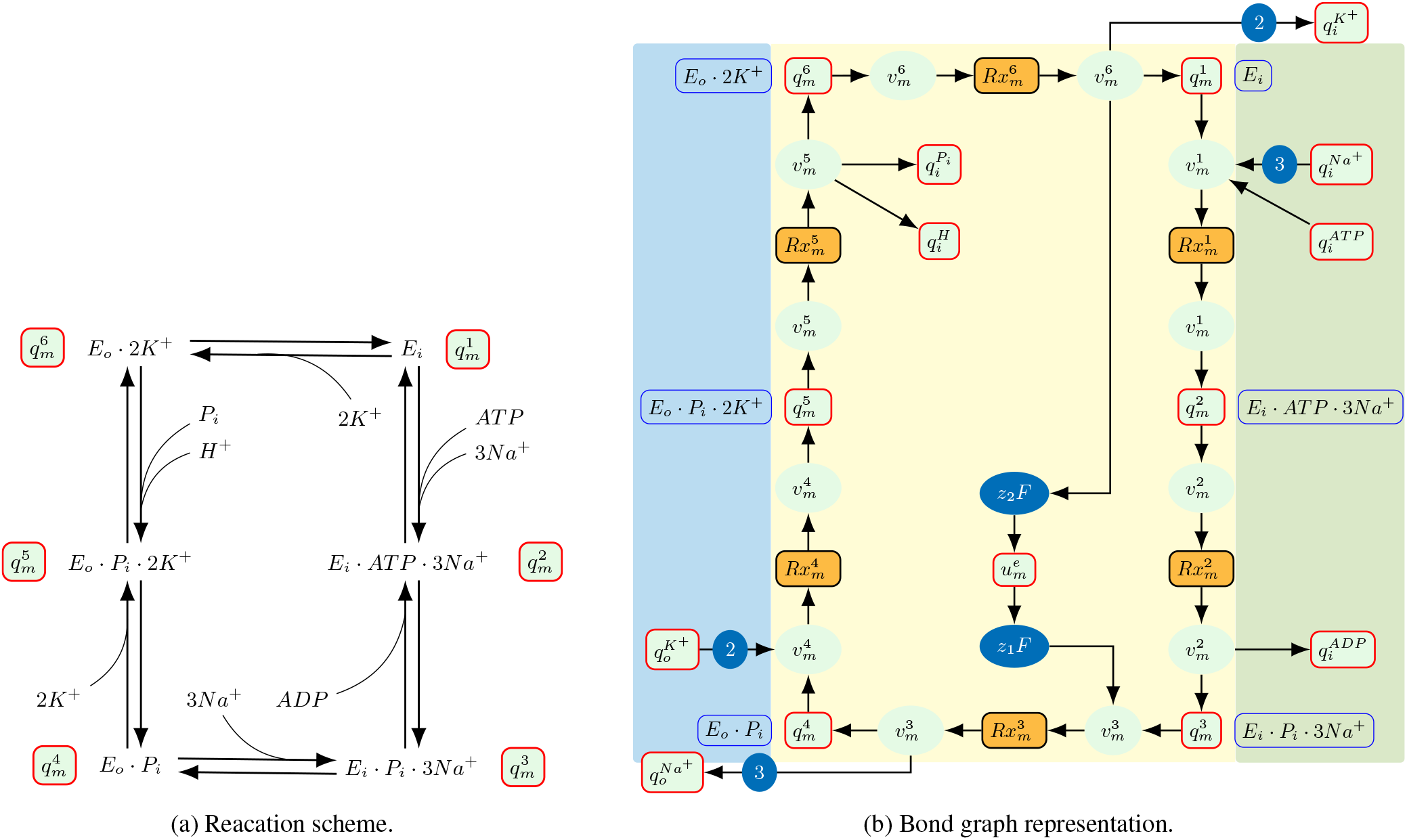
The reaction scheme and bond graph representation of the NKA. (1) The enzyme (*E*_*i*_) binds three intracellular Na^+^ ions from the cytoplasm and ATP binds to the enzyme-3Na^+^ complex, leading to (*E*_*i*_ · *AT* · *P* 3*Na*^+^); (2) The enzyme-3Na^+^-ATP complex undergoes phosphorylation and ADP is released, leading to the formation of the enzyme-3Na^+^-P complex (*E*_*i*_ · *P*_*i*_ · 3*Na*^+^); (3) The enzyme-3Na^+^-P complex undergoes a conformational change to the exoplasmic side-open state, releasing three Na^+^ ions to the extracellular space (*E*_*o*_ · *P*_*i*_); (4) Two extracellular K^+^ ions bind to the enzyme in the exoplasmic side-open state, leading to enzyme-2K^+^ complex (*E*_*o*_ · *P*_*i*_ · 2*K*^+^); (5) The enzyme-2K^+^ complex (*E*_*o*_ · *P*_*i*_ · 2*K*^+^) undergoes dephosphorylation, releasing *P*_*i*_ and *H*^+^, ending up with the enzyme-2K^+^ complex (*E*_*o*_ · 2*K*^+^); (6) The enzyme-2K^+^ complex (*E*_*o*_ · 2*K*^+^) undergoes a conformational change to the cytoplasmic side-open state (*E*_*i*_), leading to the release of two K^+^ ions into the cytoplasm.

### Energetic analysis of NKA models

The NKA extrudes three Na^+^ ions from the intracellular space and imports two K^+^ ions into the cell per molecule of ATP hydrolyzed. The overall reaction can be summarised as:

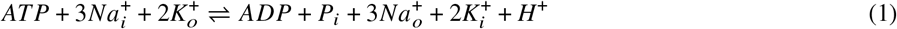

where the subscripts *i* and *o* denote intracellular and extracellular species, respectively.

The ATP hydrolysis reaction is exergonic under physiological conditions, releasing free energy that drives the active transport of Na^+^ and K^+^ ions against their electrochemical gradients. The chemical reaction for ATP hydrolysis is given by:

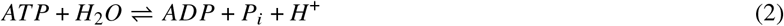

The Gibbs free energy change associated with ATP hydrolysis can be expressed as:

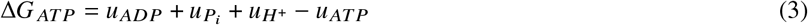

where *u*_*ATP*_, *u*_*ADP*_, 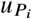, and 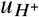 are the chemical potentials of ATP, ADP, inorganic phosphate (Pi), and protons *H*^+^, respectively.

The thermodynamic efficiency of the NKA at steady state can be estimated using Equation 4 (13) below:

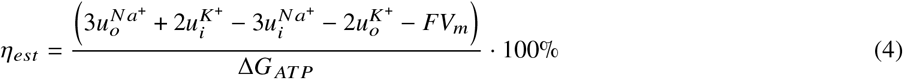

where 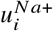 and 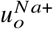 are the chemical potentials of intracellular and extracellular Na^+^, respectively, 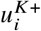 and 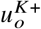 are the chemical potentials of intracellular and extracellular K^+^, respectively, *F* is Faraday’s constant (96485 C mol^-1^), and *V*_*m*_ is the membrane potential.

The Gibbs free energy change of the overall NKA reaction can be calculated as:

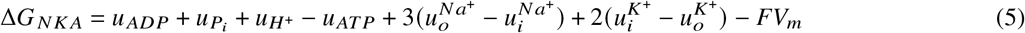

Under physiological conditions, the membrane potential, particularly for cardiac cells, varies over time and this variation affects the thermodynamic efficiency of the NKA (13, 49).

The energy flows in the NKA are shown in Figure 3. ATP hydrolysis releases energy to drive the transport of Na^+^ and K^+^ ions. Some energy is dissipated as heat, while the rest is stored as chemical energy in the concentration gradients of Na^+^ and K^+^ across the cell membrane and electrical energy in the membrane potential.

**Figure 3:**
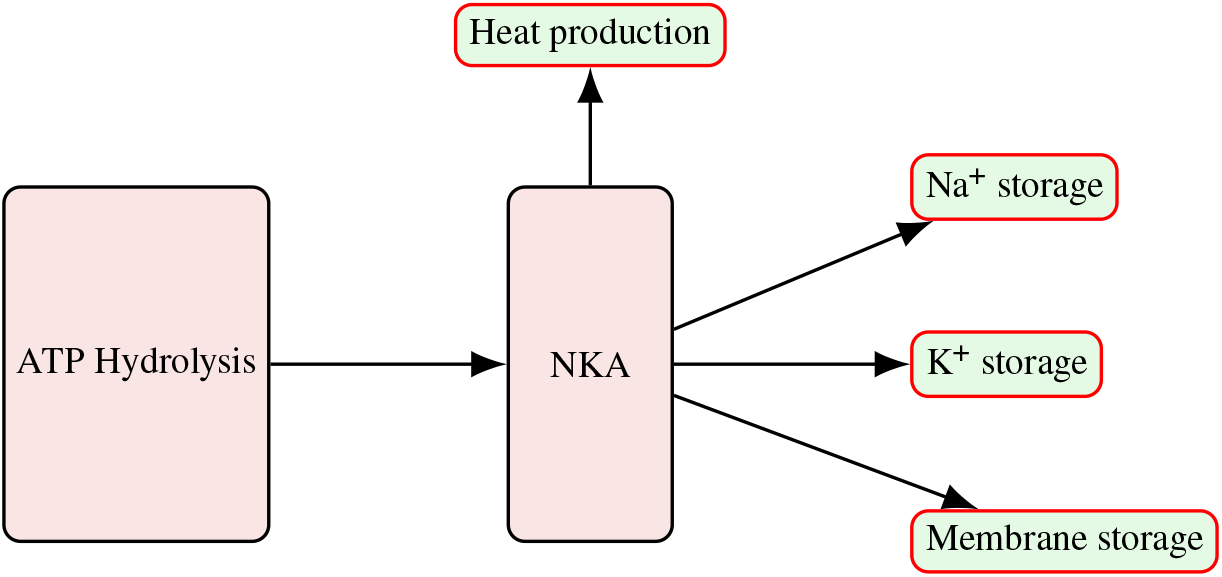
Schematic diagram of energy flows through the NKA.

In this study, we perform a detailed energetic analysis of the NKA models using the bond graph framework to account for the time-varying nature of the membrane potential and other dynamic factors. The total energy supplied by ATP hydrolysis over a time interval *t* can be calculated as:

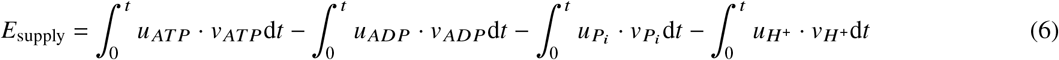

where *v*_*ATP*_, *v*_*ADP*_, 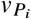, and 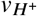 are the flow rates of ATP hydrolysis, ADP production, Pi production, and proton production, respectively.

The energy dissipated by the NKA depends on the model structure, and can be calculated as:

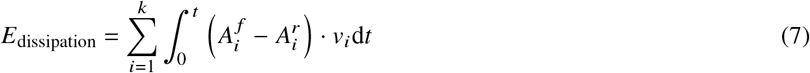

where 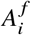 and 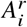 are the forward and reverse affinities of reaction *i*, respectively, *v*_*i*_ is the flow rate of reaction *i*, and *k* is the total number of reactions in the NKA models. For the 15-state model, *k* = 15, and for the 6-state model, *k* = 6.

The chemical energy stored in the concentration gradients of Na^+^ and K^+^ concentration gradients across the cell membrane can be calculated as:

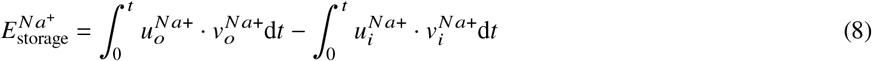

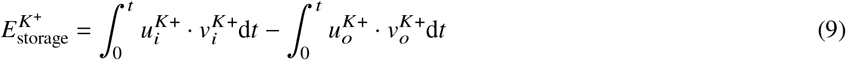

where 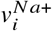 and 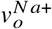 are the flow rates of intracellular and extracellular Na^+^, respectively; 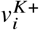 and 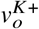 are the flow rates of intracellular and extracellular K^+^, respectively.

The total stored chemical energy can then be calculated as:

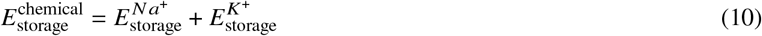

The electrical energy stored in the membrane potential can be calculated as:

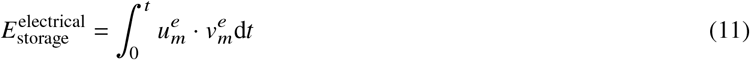

where *V*_*m*_ is the membrane potential and *i*_*m*_ is the current mediated by the NKA.

The total energy balance of the NKA model can be written as:

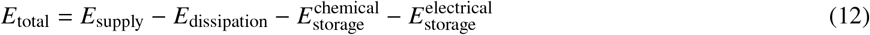

where *E*_total_ should be zero to satisfy the first law of thermodynamics.

The thermodynamic efficiency of the NKA can be defined as the ratio of the useful energy stored to the total energy supplied:

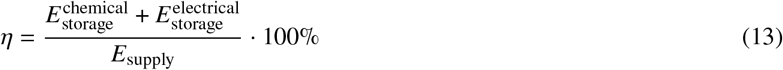

The efficiency defined in Equation 13 is different from the efficiency estimated by Equation 4, which is based on the steady state assumption and does not account for the time-varying nature of the membrane potential and other dynamic factors. The efficiency defined in Equation 13 provides a more comprehensive and accurate measure of the energetic efficiency of the NKA under dynamic conditions. This efficiency can be used to assess the performance of the NKA under different physiological and pathophysiological conditions, and to compare the energetic efficiency of different NKA models.

Please note that the normal range of NKA efficiency is between 0 and 100%, where the ATP hydrolysis is the only source of energy. When the calculated efficiency is greater than 100%, it indicates that there are other sources of energy contributing to the transport process, such as the energy stored in the concentration gradients of Na^+^ and K^+^ or the electrical energy stored in the membrane potential. In such cases, the efficiency can be interpreted as the ratio of useful energy stored to the total energy supplied from ATP hydrolysis, and it may exceed 100% due to the additional energy contributions from other sources. At extreme conditions, the calculated efficiency can be negative, which indicates that the NKA operates in reverse mode under certain conditions, using the energy stored in the concentration gradients of Na^+^ and K^+^ or the electrical energy stored in the membrane potential to drive the synthesis of ATP. The reversal of the pump cancels out some of the forward fluxes, which results in reduced efficiency. When the accumulated *E*_supply_ from the reverse fluxes is greater than the accumulated value from the forward fluxes, the value of *E*_supply_ becomes negative,and so does the efficiency.

The chemical energy conversion efficiency of the NKA can be defined as the ratio of chemical energy stored to the total energy supplied:

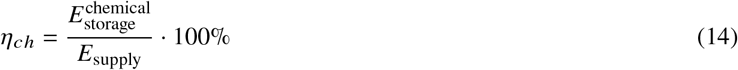

The electrical energy conversion efficiency of the NKA can be defined as the ratio of electrical energy stored to the total energy supplied:

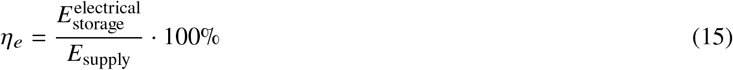

The ratio of heat production (i.e. energy dissipation) can also be calculated to assess the energy loss in the NKA:

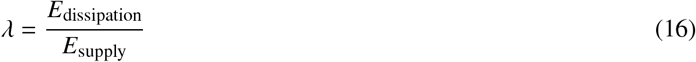

The ratio of chemical energy stored to electrical energy stored can be calculated to assess the distribution of energy storage in the NKA:

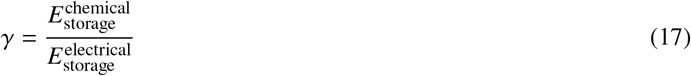

### Simulation conditions

In this paper, we focus on the energetic analysis of NKA under physiological conditions for cardiac cells, and we set up the simulation conditions based on the literature review as follows. Since ATP hydrolysis serves as the energy source for the NKA, the free energy change of ATP hydrolysis (Δ*G*_*ATP*_) plays a crucial role in determining the activity of the NKA. The Δ*G*_*ATP*_ depends on the concentrations of ATP, ADP, Pi, and *H*^+^ (i.e., pH value).

The intracellular ATP concentration ranges from 1.92–7.47 mM with an average of 4.4 mM (50), while the recent study showed lower diastolic cytosolic ATP levels around 457 ± 47 *μ*M (less than 1 mM), which is less than previously estimated, and the ATP concentration fluctuates during excitation-contraction coupling (51).

The cytosolic ADP concentrations of in situ mouse, rat and guinea pig heart are 13, 18 and 22 *μ*M (52), respectively, while the extrapolated value in human heart is 45.66 *μ*M for a body weight of 70 kg. The experimental observation of Pi in the heart in vivo has not been studied yet, while the baseline Pi of a model prediction in the canine myocyte cytoplasm is around 0.29 mM and increases with work rate in heart cells (53).

The simulation in (54) reported ATP, ADP, Pi and pH values of 5.61 mM, 25.4 *μ*M, 1.16 mM and 7.15, respectively. In general, the physiological intracellular pH ranges from 7.15 to 7.25 (55).

Intracellular Na^+^ concentration is normally from 4 to 16 mM (56), while the level can rise by 3 mM above this value in heart failure conditions (57). This increase is due to increased Na^+^ influx while the NKA function is unchanged (57). Around 98% of potassium (K^+^) resides in the cytosol at 140–150 mM and the extracellular K^+^ concentration ranges from 3.5 to 5 mM (58).

We use the simulation conditions as shown in Table 1 to examine the distribution of energy flows in different NKA models. In addition, we vary one parameter at a time to investigate its effect on the function and energetics of the NKA and the values used are shown in the Results section.

**Table 1:**
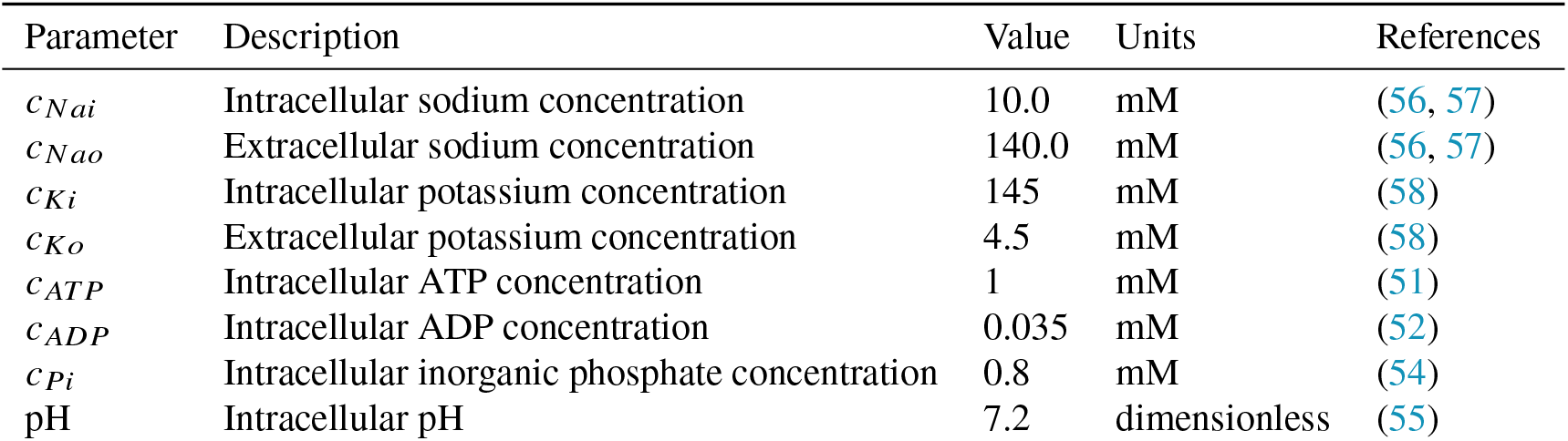
Default parameters for NKA.

| Parameter | Description | Value | Units | References |
| --- | --- | --- | --- | --- |
| $c_{\text{Nai}}$ | Intracellular sodium concentration | 10.0 | mM | (56, 57) |
| $c_{\text{Nao}}$ | Extracellular sodium concentration | 140.0 | mM | (56, 57) |
| $c_{\text{Ki}}$ | Intracellular potassium concentration | 145 | mM | (58) |
| $c_{\text{Ko}}$ | Extracellular potassium concentration | 4.5 | mM | (58) |
| $c_{\text{ATP}}$ | Intracellular ATP concentration | 1 | mM | (51) |
| $c_{\text{ADP}}$ | Intracellular ADP concentration | 0.035 | mM | (52) |
| $c_{\text{Pi}}$ | Intracellular inorganic phosphate concentration | 0.8 | mM | (54) |
| pH | Intracellular pH | 7.2 | dimensionless | (55) |

## RESULTS

### Fit 6-state NKA models to experimental data

We applied the experimental conditions in (30) to fit the 6-state bond graph model to the experimental data using the differential evolution method in SciPy (59), shown in Figure 4.

**Figure 4:**
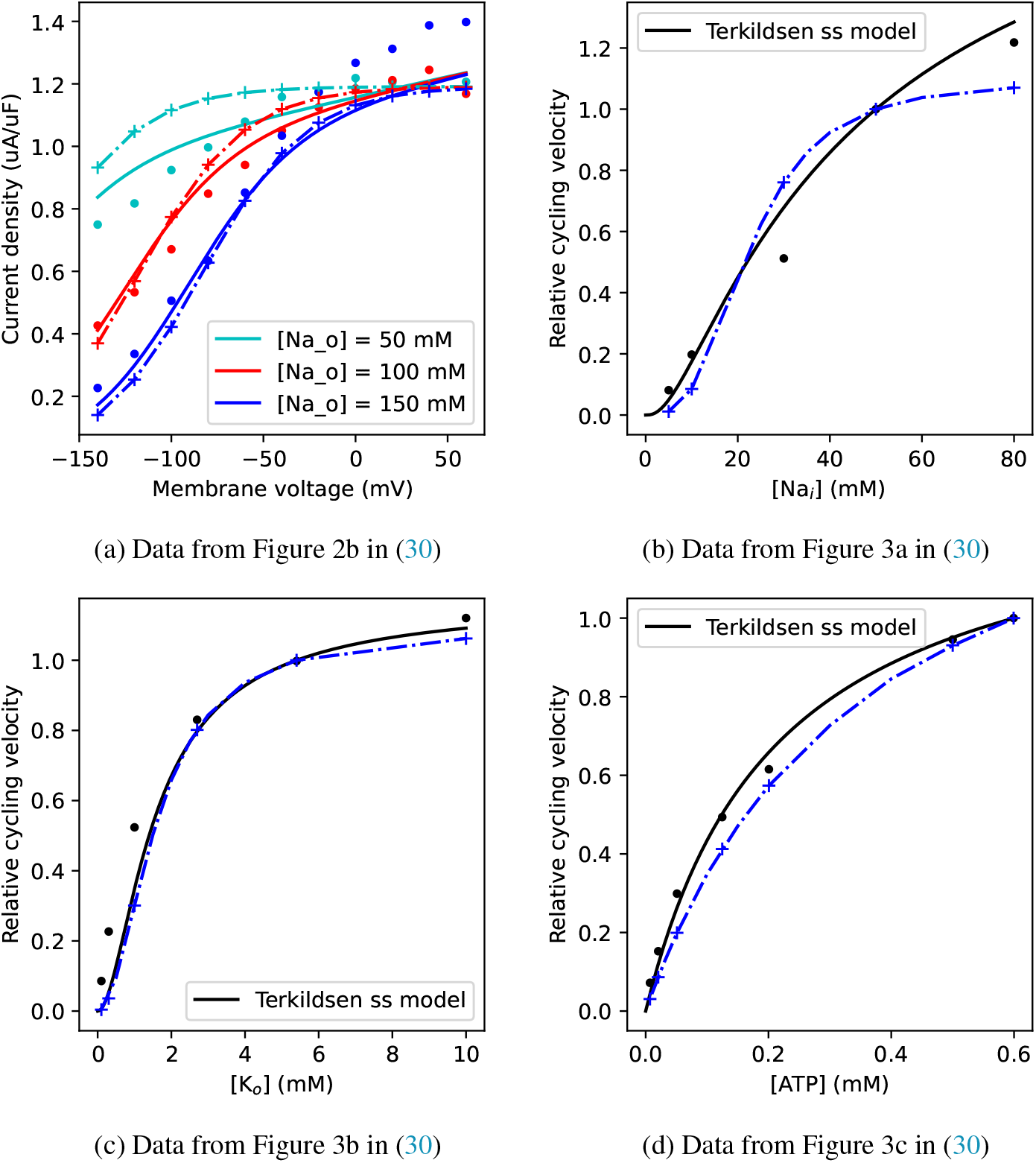
Fit of NKA models to experimental data and Terkildsen model after correction in (30). The solid lines are simulated data from the kinetic model in (30), while the solid dots are experimental data. The dashed lines are simulated data from the steady-state 6-state bond graph model, and the ‘+’ symbols are produced by the full 6-state bond graph model. (a) NKA current-voltage relationship at different extracellular Na^+^ concentrations; (b) NKA current as a function of intracellular Na^+^ concentration; (c) NKA current as a function of extracellular K^+^ concentration; (d) NKA current as a function of ATP concentration. The simulation conditions are the same as in (30).

The simulated data are comparable to the kinetic model in (30) and experimental data under most conditions, except for the low extracellular Na^+^ concentration (50 mM) in Figure 4 (a). The 6-state model has higher current density when the extracellular Na^+^ concentration is 50 mM compared to the kinetic model in (30), while extracellular Na^+^ concentration is normally around 140 mM (56) under physiological conditions. In the following sections, we will focus on the comparison of the 15-state and 6-state bond graph models with regard to thermodynamic aspects.

### Energy flows through NKA

We applied a series of action potentials (with baseline cycle length (BCL) of 1000 ms) to the NKA models while keeping the Na^+^ and K^+^ concentrations constant. The action potentials were simulated using the Faber-Rudy model (60) with corrected T type calcium current (61), which was retrieved from PMR (https://models.physiomeproject.org/e/260). The other simulation conditions are shown in Table 1. We then calculated the energy flows through the NKA over ten cycles of the action potential. We plotted the results in Figure 5 (a and b).

**Figure 5:**
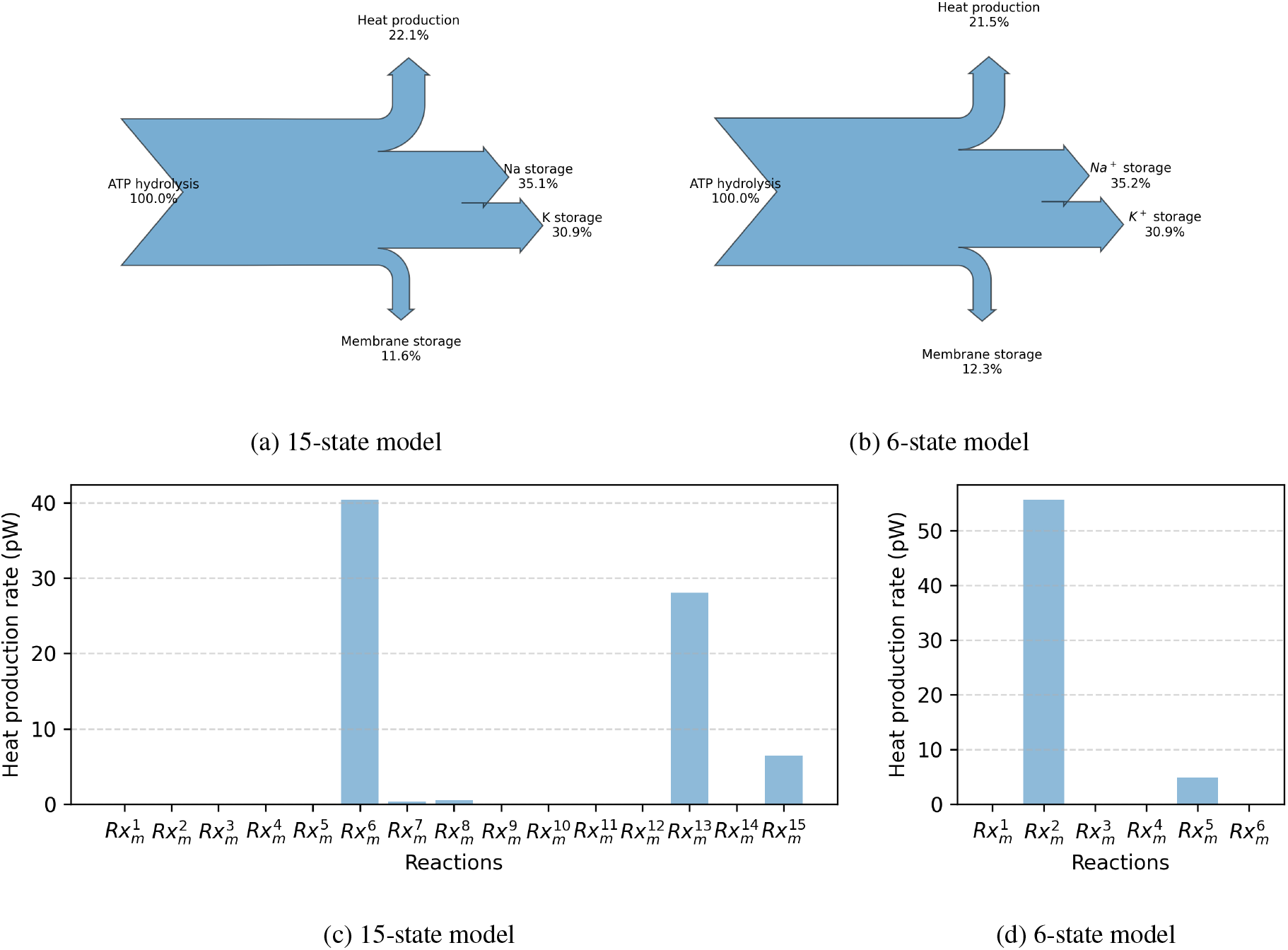
Energy flow diagrams and heat production rates of (a) 15-state model and (b) 6-state model of NKA under physiological conditions.

**Figure 6:**
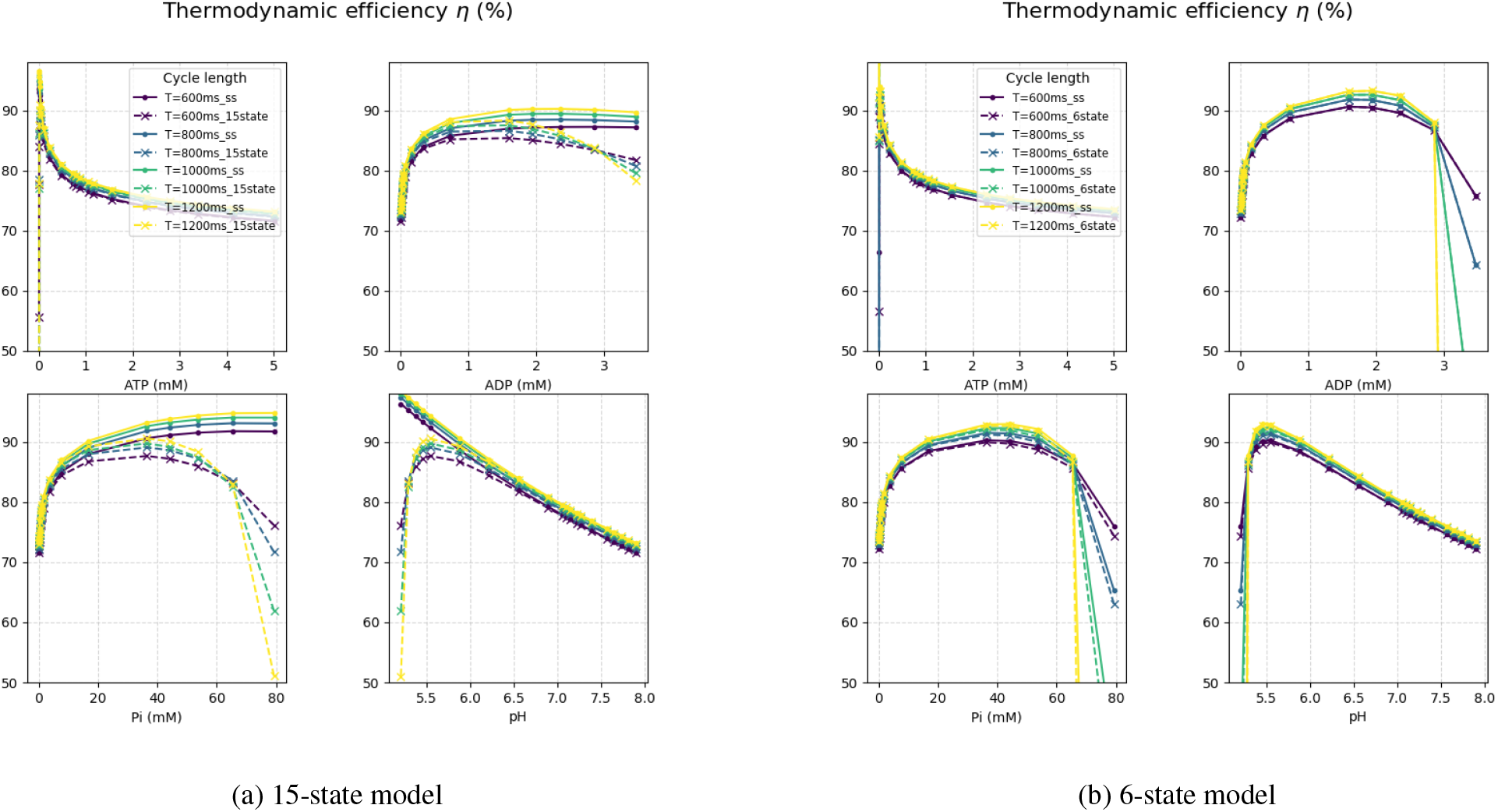
Thermodynamic efficiency of NKA (a) (15-state model) and (b) (6-state model). The right y-axis shows the corresponding Δ*G*_*ATP*_ values, indicated by the blue dots in the plots.

We observe that around 22% of the energy supplied by ATP hydrolysis is dissipated as heat, 12% is stored as electrical energy, and 66% is stored as chemical energy in both models under physiological conditions. The thermodynamic efficiency of both models is around 78%, which is consistent with previous theoretical studies (13, 49). We also calculated the heat production rates of both models, shown in Figure 5 (c and d). In the 15-state model, the energy dissipation mainly occurs during the reaction steps of 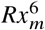 when ADP is released, and 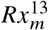 when *P*_*i*_ and *H*^+^ are released and 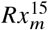 when the enzyme is translocated from extracellular to intracellular after ATP binding. In the 6-state model, the energy dissipation mainly occurs during the reaction steps of 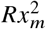 when ADP is released, and 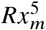 when *P*_*i*_ and *H*^+^ are released.

According to the theoretical study in (13), the NKA is a highly efficient molecular machine due to the micro conformational changes that minimize energy loss during ion transport. The data in their study showed that the efficiency of NKA ranges from around 50% to 88% depending on operating conditions. The simulation results in our study are within this range, which further supports the notion that NKA operates with high thermodynamic efficiency under physiological conditions.

Another theoretical study in (49) speculated that the ratio of Na^+^ to K^+^ transport is optimized to achieve maximum efficiency, cell volume homeostasis, and stable electrochemical gradients.

In the following section, we will further investigate the thermodynamic efficiency of the NKA under a range of conditions, which provides insights into the energy transduction mechanisms of the NKA. With this understanding, scientists can better select appropriate NKA models for their research and develop more accurate models in the future.

### Thermodynamic efficiency of NKA

To investigate the thermodynamic efficiency of the NKA under physiological conditions, we applied the action potential with different BCL to simulate the NKA models, and calculated the thermodynamic efficiency over ten cycles. We also vary the ATP, ADP, Pi concentrations and pH values to examine their effects on the thermodynamic efficiency of the NKA. When varying one parameter, the other parameters are set to the default values in Table 1. The corresponding Δ*G*_*ATP*_ values range from −46 kJ/mol to −62 kJ/mol.

With increased ATP concentrations, the thermodynamic efficiency of both models rapidly increases initially and then drops to a relatively stable level around 70%. When the ATP concentration is very low, the NKA operates in reverse mode, using the energy stored in the concentration gradients of Na^+^ and K^+^ or the electrical energy stored in the membrane potential to drive the synthesis of ATP, which results in negative efficiency. As the ATP concentration increases, the NKA starts to hydrolyze ATP and transport ions, leading to a rapid increase in efficiency. However, at very high ATP concentrations, the efficiency drops slightly due to increased energy dissipation as heat.

With increased ADP concentrations and Pi concentrations, the thermodynamic efficiency of both models increases rapidly at low concentrations and then reaches a plateau. Afterward, the full 15-state bond graph models present a decrease in efficiency at high concentrations, while the steady-state models maintain relatively stable efficiency. The 6-state full bond graph model and its steady-state model show similar trends under most conditions. The drop in efficiency at high ADP and Pi concentrations is due to the NKA synthesizing ATP instead of hydrolysing ATP.

The study (53) showed the free ADP slightly increases from 0.042 mM to 0.064 mM, while the Pi concentration is approximately 0.29 mM and increases to 2.3 mM at high work rate. According to our simulation, the increased ADP and Pi concentrations during high work rate will make the NKA operate at a higher efficiency.

In the majority of the pH range (5.5-8), the thermodynamic efficiency of both models decreases linearly with increased pH values.

From the figures, we can see that the ATP, ADP, Pi concentrations and pH values affect the thermodynamic efficiency of NKA in a different manner even though the Δ*G*_*ATP*_ values are within the same range. To further investigate the effect of Δ*G*_*ATP*_ on the thermodynamic efficiency of NKA, we plotted the thermodynamic efficiency of both models under different Δ*G*_*ATP*_ values by varying the ATP, ADP, Pi concentrations and pH values, shown in Figure 7.

**Figure 7:**
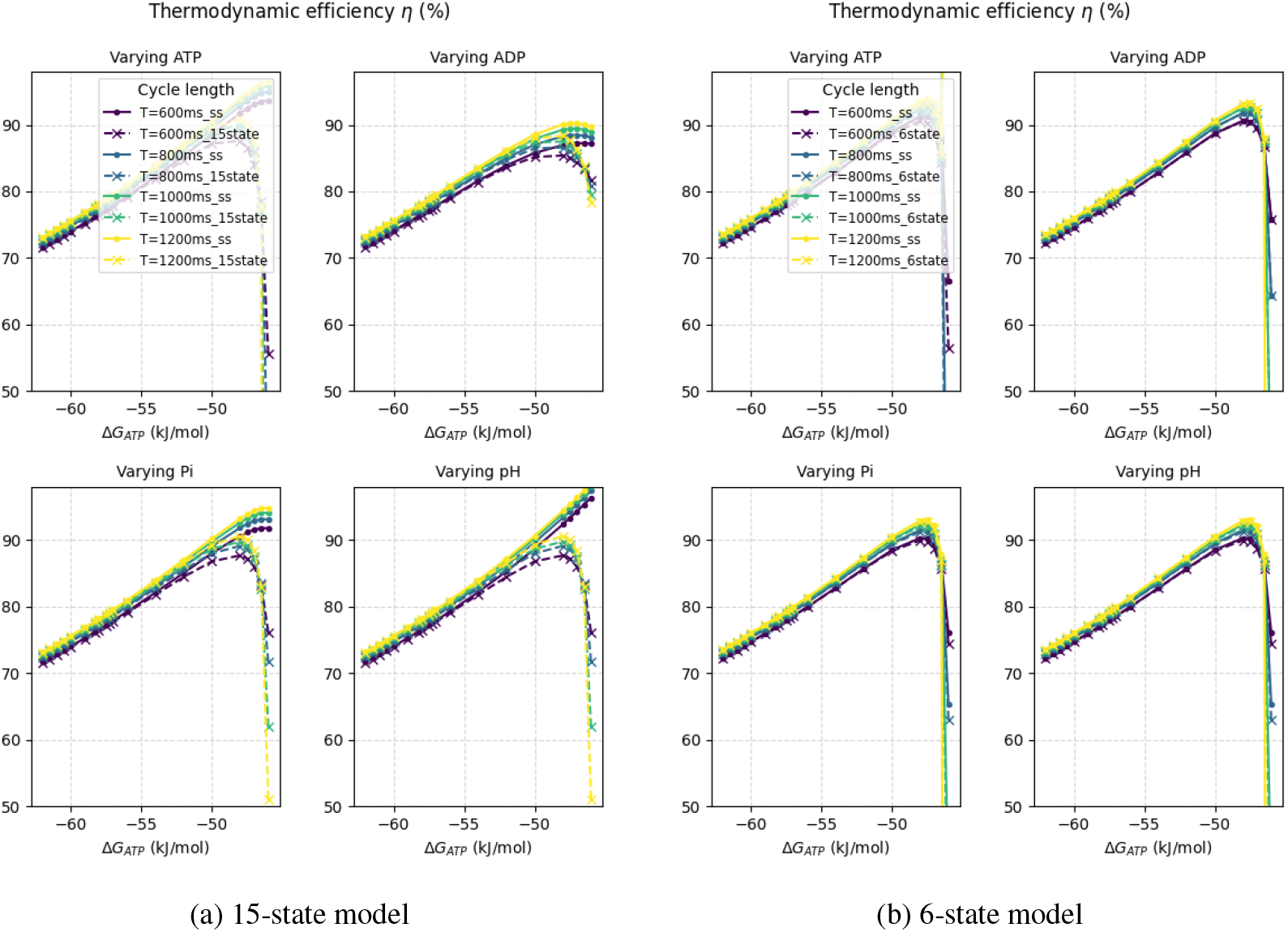
Thermodynamic efficiency of NKA (a) (15-state model) and (b) (6-state model) under different Δ*G*_*ATP*_ values.

We can see that the thermodynamic efficiency of both models increases with lower Δ*G*_*ATP*_ values (less negative) until it drops in the full bond graph models after the Δ*G*_*ATP*_ reaches around −48 kJ/mol. We calculated the combined free energy for the chemical and electrical stores at resting potential (85 mV) Δ*G*_*NKA*_ using Equation 5, which becomes positive when Δ*G*_*ATP*_ = −46 kJ/mol. This seems aligned with the study (62, 63). Kammermeier estimated that the NKA in the heart requires a minimum Δ*G*_*ATP*_ of −46 kJ/mol, while the study (63) found that when Δ*G*_*ATP*_ was approximately −49 kJ/mol, little activity of NKA at membrane potentials more negative than −75 mV was observed in sheep Purkinje cells.

Since Equation 5 does not account for the dynamics of the NKA, it cannot explain the differences between the 15-state and 6-state models or the drop at different BCL values. To investigate the mechanism behind the drop in thermodynamic efficiency at low Δ*G*_*ATP*_ values, we computed the biochemical conversion efficiency *η*_*ch*_ and electrical conversion efficiency *η*_*e*_ using equations 14 and 15, respectively. Both *η*_*ch*_ and *η*_*e*_ contribute to the overall thermodynamic efficiency drop, while the BCL of the action potential has a more obvious effect on *η*_*e*_, as shown in Figure 8.

**Figure 8:**
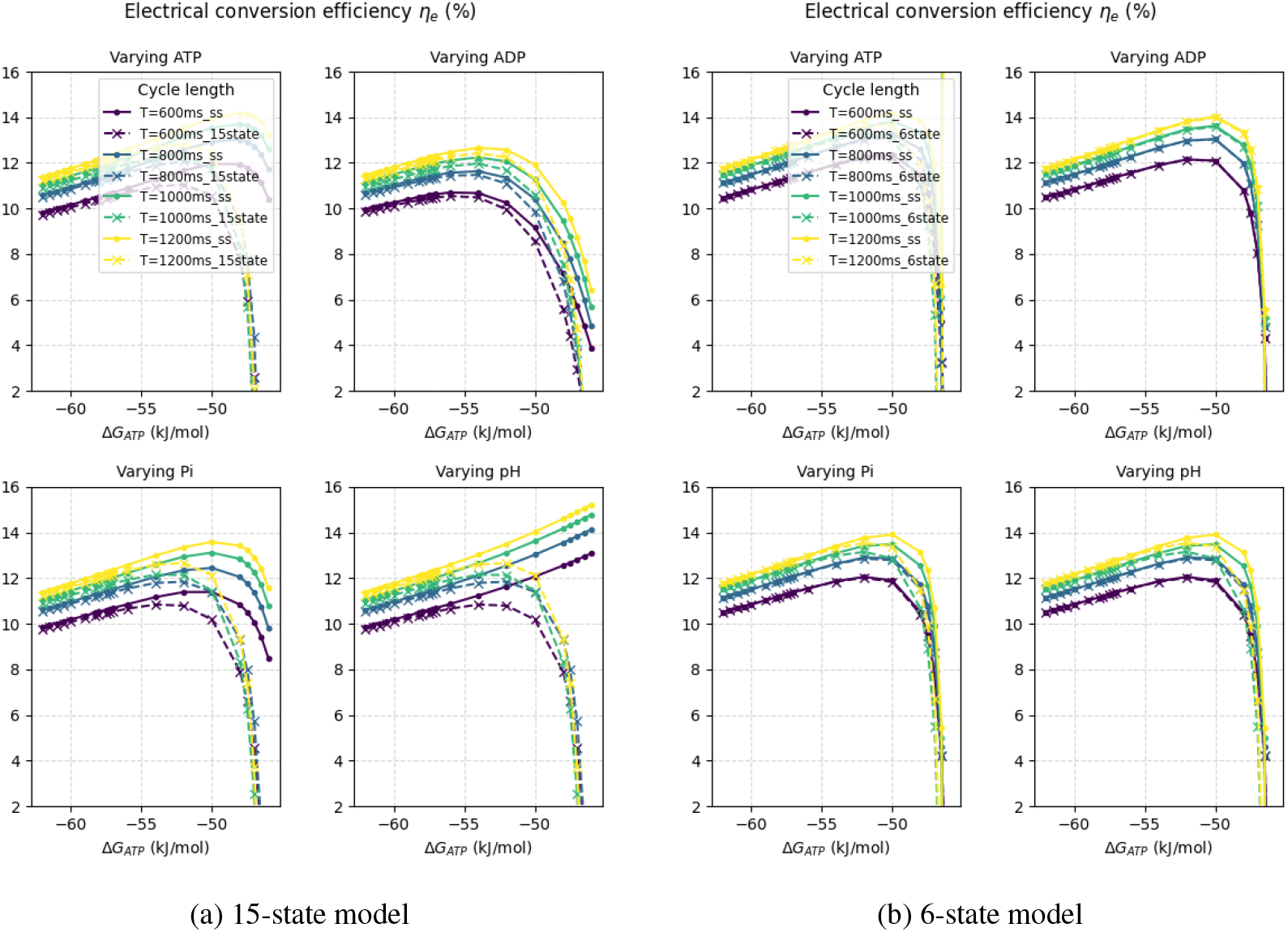
Electrical conversion efficiency of NKA (a) (15-state model) and (b) (6-state model) under different Δ*G*_*ATP*_ values.

We can see that the electrical conversion efficiency of both models slightly increases with lower Δ*G*_*ATP*_ values (less negative), and there is a drop to negative in full bond graph models after the Δ*G*_*ATP*_ reaches around −48 kJ/mol, which aligns with the drop in thermodynamic efficiency.

The trends of the electrical conversion efficiency of both models are similar under varying ATP, Pi concentrations, but differ when ADP and pH values are varied. With varied ADP, the electrical conversion efficiency of the 15-state model starts to decrease earlier than that of the 6-state model; with varied pH, the electrical conversion efficiency of the 15-state steady-state model does not drop at all. Theese differences may be due to the different confirguration of multiple reaction steps of the two models.

Under physiological conditions, the current mediated by the NKA is outward. When the free energy available from ATP hydrolysis (Δ*G*_*ATP*_) decreases (less negative), the current reduces and eventually reverses direction, leading to an inward current, as observed experimentally in (63). In our simulation, the energy flow (i.e., power output) of the membrane potential becomes negative when Pi or ADP concentration is increased, or pH is decreased, which means that the charges over the membrane act as energy source. This leads to a drop in electrical conversion efficiency and consequently a drop in thermodynamic efficiency of the NKA.

It is worth noting that the electrical conversion efficiency is lower at higher work rate (i.e. shorter BCL).

### Inhibition of NKA by reduced Δ*G*_*ATP*_

Jansen et al. (64) investigated the effect of reduced free energy available from ATP hydrolysis on NKA activity in an oxygenated isolated rat heart. In their study, they could not distinguish whether the effect on NKA activity was due to changes in ATP, ADP, Pi, or pH, but they observed that as Δ*G*_*ATP*_ decreased, there was a reduction in *Rb*^+^ uptake, indicating reduced NKA activity.

We ploted the mean turnover rate of NKA under different Δ*G*_*ATP*_ values by varying the ATP, ADP, Pi concentrations and pH values, shown in Figure 9.

**Figure 9:**
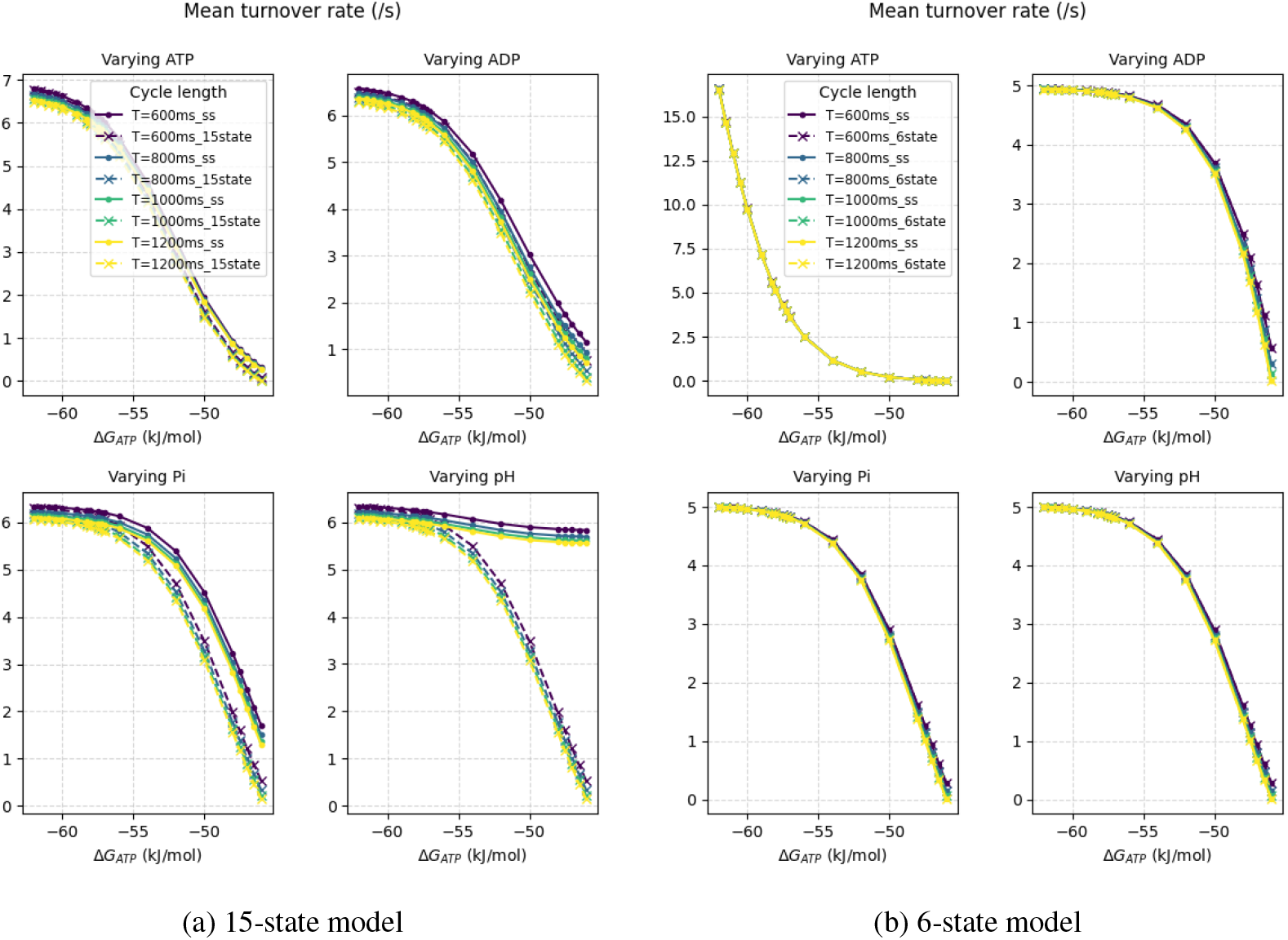
Mean flow rate of NKA (a) (15-state model) and (b) (6-state model) under different Δ*G*_*ATP*_ values.

We can see that the mean turnover rate of both models decreases with lower Δ*G*_*ATP*_ values (less negative), which aligns with the experimental observation in (64).

Huang et al. (65) found that the inhibition of NKA activity by Pi depended on the pH range, with greater inhibition at lower pH values of 6.1–7, but less inhibition with increased pH of 7–8.5. With 5 mM Pi, the NKA activity was inhibited by around 30% at pH 7. In our simulation, we can see that the inhibition of NKA activity by Pi is more pronounced at lower Δ*G*_*ATP*_ values, which is associated with lower pH values and higher Pi concentrations.

Increases in intracellular Pi and decreases in intracellular pH and ATP were associated with ischemic conditions (53, 65), and these changes are associated with the inhibition of NKA activity if other conditions are kept constant according to our simulation.

To understand how the inhibition of NKA activity affects its thermodynamic efficiency, we plotted the thermodynamic efficiency of both models under different mean turnover rates by varying the ATP, ADP, Pi concentrations and pH values, shown in Figure 10.

**Figure 10:**
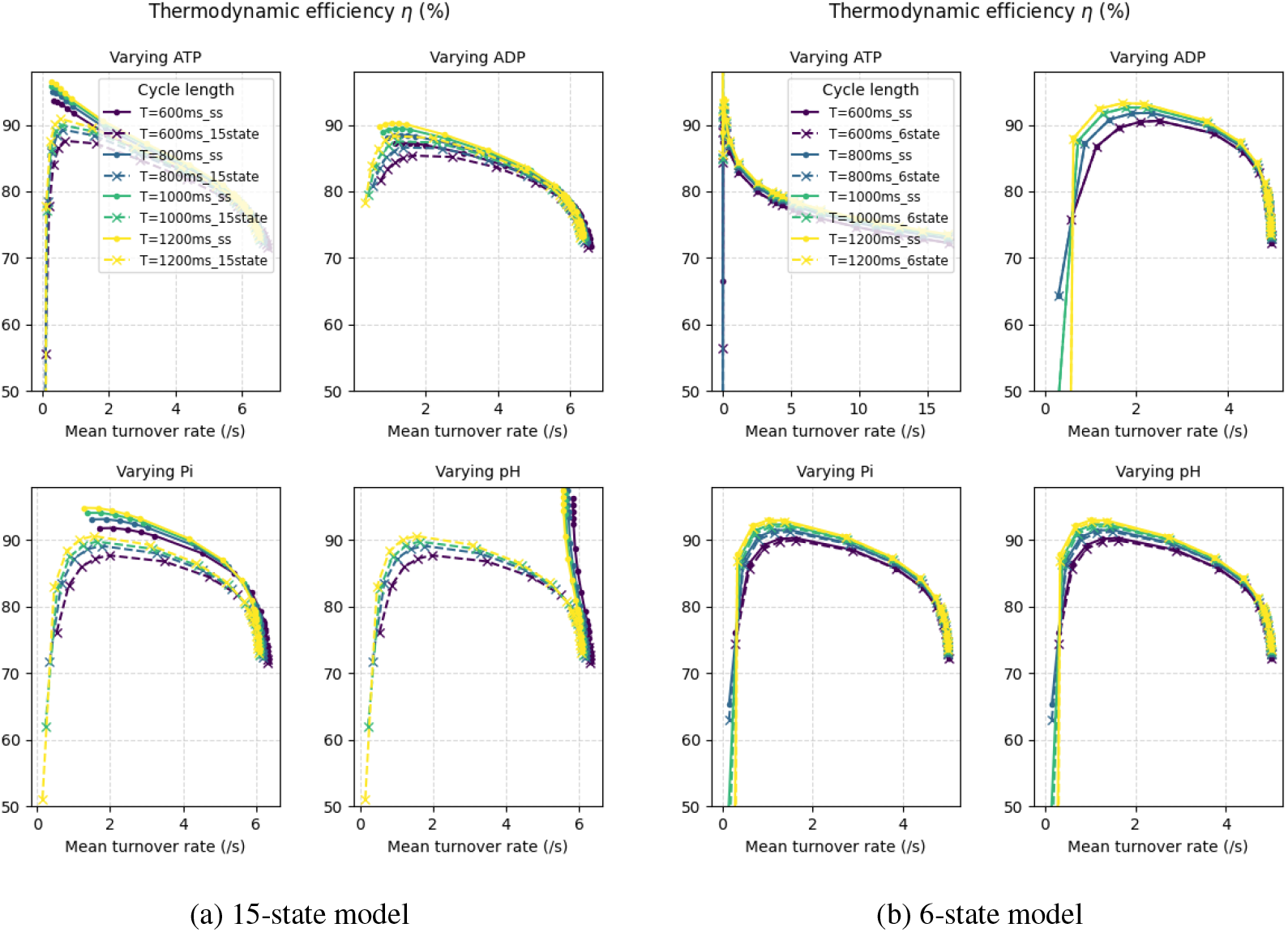
Thermodynamic efficiency of NKA (a) (15-state model) and (b) (6-state model) under different mean turnover rates.

We can see that the thermodynamic efficiency of both models increases with reduced mean turnover rates, which aligns with the study in (13). The drop in thermodynamic efficiency at very low mean turnover rates in full bond graph models is due to the drop in electrical conversion efficiency, as discussed in the previous section.

### Response of NKA to changes in intracellular Na^+^ concentration and extracellular K^+^ concentration

The reduction of free energy available from ATP hydrolysis results in an accumulation of intracellular Na^+^ ions (64). The increase was noted when Δ*G*_*ATP*_ fell below −52 kJ/mol and larger increases were noted when Δ*G*_*ATP*_ dropped below −48 kJ/mol. This finding agreed with the experimental observation that little activity of NKA at membrane potentials more negative than −75 *mV* was observed when Δ*G*_*ATP*_ was approximately −49 kJ/mol (63). These observations are consistent with our simulation results, shown in Figure 9, where the decrease in mean turnover rate of both models is more significant at lower Δ*G*_*ATP*_ values (less negative).

Despa et al. (57) reported that the resting intracellular Na^+^ concentration is higher in heart failure conditions due to increased Na^+^ influx while the NKA function is unchanged. They also reported increased intracellular Na^+^ during stimulation in rabbit ventricular myocytes. Nelson et al.(66) reported increasing stimulation rate from 60 beats per minute (bpm) to 180 bpm led to a transient Na^+^ peak followed by a lower Na^+^ level in both rat and human ventricular myocardium. As the cardiac action potential is initiated by Na^+^ influx, the increased stimulation rate leads to increased Na^+^ influx. The peak is due to the initial accumulation of intracellular Na^+^ ions, while the subsequent lower Na^+^ level is due to the increased NKA activity to extrude more Na^+^ ions.

A simulation study (67) showed that extracellular K^+^ increased during acute myocardial ischemia due to the combined effect of activation of ATP-dependent K^+^ current, an ischemic Na^+^ influx and inhibition of NKA.

To investigate the response of NKA to changes in intracellular Na^+^ concentration and extracellular K^+^ concentration when Δ*G*_*ATP*_ remains constant, we varied the intracellular Na^+^ concentration from 4 mM to 16 mM or extracellular K^+^ concentration from 3.5 mM to 5.5 mM, shown in Figure 11.

**Figure 11:**
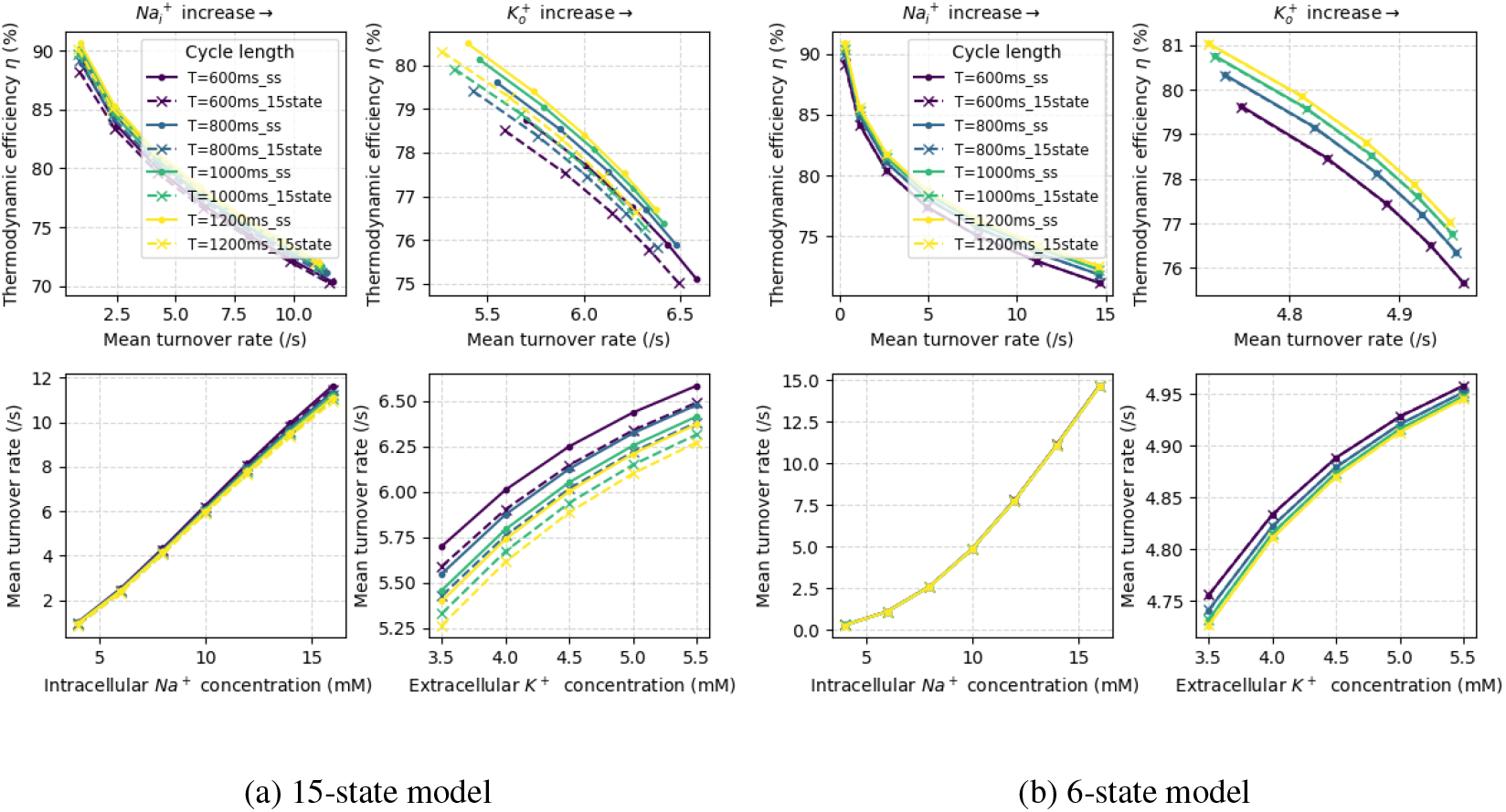
Thermodynamic efficiency and mean turnover rate of NKA (a) (15-state model) and (b) (6-state model) under different intracellular Na^+^ and extracellular K^+^ concentrations. The left panels of (a) and (b) show the thermodynamic efficiency of NKA with increased mean turnover rate due to increased intracellular Na^+^ concentration, while the right panels of (a) and (b) show the thermodynamic efficiency of NKA with increased mean turnover rate due to decreased extracellular K^+^ concentration.

We can see that the mean turnover rate of both models increases with increased intracellular Na^+^ concentration or extracellular K^+^ concentration. This confirms that the NKA activity is regulated by intracellular Na^+^ concentration and extracellular K^+^ concentration (57, 66) provided Δ*G*_*ATP*_ remains constant. We note that constant Δ*G*_*ATP*_ does not imply constant energy release from ATP hydrolysis. Instead, the power output from ATP hydrolysis increases with increased intracellular Na^+^ concentration or extracellular K^+^ concentration because of increased NKA activity. That is, more energy is needed to extrude more Na^+^ ions and import more K^+^ ions against their electrochemical gradients. As shown in the study (66), the NKA can respond to increased intracellular Na^+^ concentration during stimulation by increasing its activity to maintain ion homeostasis provided that sufficient energy is available from ATP hydrolysis.

The thermodynamic efficiency of both models decreases with increased turnover rate, which aligns with the study in (13). This suggests that more energy is dissipated as heat at higher NKA activity.

## DISCUSSION

A number of kinetic models of NKA have been proposed to investigate its function and regulation under various conditions (5, 25–29). While these models follow the Albers–Post scheme (2, 3), they differ in the number of conformational states and the level of detail in describing the conformational changes and ion binding/unbinding steps. Although cryo-electron microscopy (cryo-EM) structural studies (6–12) have provided insights into the conformational states of NKA, the detailed kinetic mechanisms of the NKA are still not fully understood. Due to the varied configuration of the reaction steps in different models, it is difficult to compare the dynamic behavior of these models apart from their steady-state kinetics. The dynamic response of the NKA is important to understand its function under physiological and pathophysiological conditions, especially in the cardiac context where the NKA plays a critical role in rhythmic cardiac excitability and contractility.

In this study, we have applied the bond graph modeling approach to investigate the energetics of NKA under various conditions in cardiac cells. Since bond graph models are thermodynamically consistent by construction, they allow us to analyze the energy flows through the NKA and calculate its thermodynamic efficiency. We can also investigate how different parameters affect the energetics of the NKA, providing insights into its energy transduction mechanisms.

The ability to compute and compare the energetics of different NKA models can help scientists select appropriate models for their research. The estimated thermodynamic efficiency at steady-state using equation 4 cannot capture the differences in dynamic behavior between models, as it does not account for the transient dynamics of the NKA. In contrast, the analysis methods (equations 6 – 17) incorporate these dynamic aspects and provide a more accurate quantification of the energetics. Our simulation results show that the full bond graph models (15-state and 6-state) present different trends in thermodynamic efficiency at low Δ*G*_*ATP*_ values compared to the steady-state models. Most cardiac action potential models, such as (60, 68), use the steady-state model of NKA, which may not capture the dynamic energetics of the NKA under certain conditions. In particular, under pathophysiological conditions such as heart failure, where the free energy available from ATP hydrolysis may be reduced, the dynamic response of the NKA and its energetic profile may differ significantly from the steady-state predictions. Therefore, for studies investigating the role of NKA under such conditions, it may be important to use a full bond graph model that captures the dynamic energetics rather than a steady-state model.

Another question is which model captures the dynamic response of NKA more accurately. Ideally, dynamic data of NKA should be used to calibrate the parameters and validate the models. However, the experimental data on the instantaneous dynamics of NKA are often limited and difficult to obtain. Since the simulated energy provides another metric to compare the models under different conditions, thermodynamic measurements under different experimental conditions can also be used to validate the models without the need for capturing the instantaneous dynamics.

Our simulation shows that the work rates affect the electrical conversion efficiency of NKA in both 15-state and 6-state models, with lower efficiency at higher work rates when the Δ*G*_*ATP*_ is sufficiently high. However, at low Δ*G*_*ATP*_ values, the electrical conversion efficiency drops to negative regardless of the work rates. Beta-blockers are commonly used in the treatment of heart failure, which reduce the heart rate and thus increase the diastolic interval (69). If the Δ*G*_*ATP*_ is still sufficiently high, the reduced heart rate may lead to increased electrical conversion efficiency of NKA, which could potentially improve its function in maintaining ion homeostasis.

The comparison of 15-state and 6-state models here is used to illustrate the application of energetic analysis to compare different models. While Figure 4 shows the kinetic similarities at steady state, the thermodynamic analysis reveals differences in the dynamic energetics of the two models, especially at low Δ*G*_*ATP*_ values. Lower Δ*G*_*ATP*_ values usually happen under pathophysiological conditions, such as heart failure. This suggests that the thermodynamic analysis can provide additional insights into the behavior of the models that may not be apparent from kinetic analysis alone, and can help guide model selection based on the specific research questions being addressed.

The theoretical study in (13) suggested that more substeps in the transport cycle lead to higher efficiency. Our simulation results do not show a clear trend in efficiency as a function of the number of substeps. Further research is needed to explore this question and determine the optimal level of detail for modeling NKA energetics. More experimental data on the energetics of NKA, as well as molecular dynamics simulations (70), would also be beneficial to validate and refine the models. In Section *Energy flows through NKA*, we identified the reaction steps that contribute most significantly to energy dissipation in the 6-state model and the 15-state model, which could be potential targets for experimental investigation to further understand the mechanisms of energy transduction in NKA and which model is more energetically realistic.

## CONCLUSION

In this study, we have demonstrated a comprehensive energetic analysis of the NKA using bond graph modeling, which ensures thermodynamic consistency and enables detailed tracking of energy flows through the pump. Our work highlights three key findings:

First, both the 15-state and 6-state bond graph models reveal that under physiological conditions, approximately 66% of the energy from ATP hydrolysis is stored as chemical energy in ion concentration gradients, 12% is converted to electrical energy via the membrane potential, and 22% is dissipated as heat. This overall thermodynamic efficiency of ~78% is consistent with theoretical predictions (13, 49), confirming that the NKA operates as a highly efficient molecular machine.

Second, our analysis demonstrates that the thermodynamic efficiency of the NKA is sensitive to the free energy available from ATP hydrolysis (Δ*G*_*ATP*_), which depends on the concentrations of ATP, ADP, Pi, and intracellular pH. Under conditions where Δ*G*_*ATP*_ is reduced below ~-48 kJ/mol, the electrical conversion efficiency drops significantly and may even reverse, consistent with experimental observations of reduced NKA activity and accumulation of intracellular Na^+^ (63, 64, 66). This suggests that energetic profiles can serve as quantities for assessing NKA function under pathophysiological conditions.

Third, we show that the NKA activity responds dynamically to changes in intracellular Na^+^ and extracellular K^+^ concentrations, with increased turnover rates at higher Na^+^ gradients, demonstrating the pump’s role in maintaining ion homeostasis. The relationship between turnover rate and thermodynamic efficiency reveals that higher activity rates incur greater energy dissipation, highlighting a fundamental trade-off between ion transport capacity and energetic efficiency.

The bond graph framework proved invaluable for this analysis, as it simultaneously enforces conservation of mass, charge, and energy, enabling us to identify which reaction steps and energy flows contribute most significantly to the overall thermodynamic behavior of the system. By comparing the 15-state and 6-state models, we demonstrate that energetic analysis can aid in model selection and validation, guiding the choice of model complexity appropriate for specific research questions.

Future work should extend this energetic analysis to other ion pumps and transporters, investigate how genetic mutations or pharmacological interventions affect NKA energetics, and develop more energetically realistic models informed by recent cryo-EM structures (6–12). Integration of these energetic models with whole-cell and tissue-level simulations will provide deeper insights into the role of NKA in cellular energy metabolism and its dysfunction in diseases such as heart failure and stroke.

## Supporting information

Supplemental model description

## AUTHOR CONTRIBUTIONS

W.A., M.P., P.J.H. and D.P.N. conceptualized and designed the research. W.A. implemented the model and analyzed the data. D.P.N. tested the reproducibility of the results. All authors contributed to the manuscript and approved the submitted version.

## ACKNOWLEDGMENTS

This project is supported by a grant from the NZ Government’s MBIE Catalyst Fund (the Biotech Digital Twin Research Programme).

## SUPPLEMENTARY MATERIAL

An online supplement to this article can be found by visiting BJ Online at http://www.biophysj.org.

