## Supplemental model description for "Energetic analysis of Na^+^/K^+^-ATPase using bond graphs"

### INTRODUCTION

This document includes the detailed model description of the Sodium-potassium ATPase pump (NKA) used in the main paper.

### METHODS

#### Bond graph model of NKA with 15 states

The reaction scheme can be found in the paper (1), while we redraw the bond graph in Figure 1 using the new notation.

The equations for conservation of mass are expressed as below.

$$\frac{dq_m^1}{dt} = v_m^{15} - v_m^1 \qquad \frac{dq_m^2}{dt} = v_m^1 - v_m^2 \qquad \frac{dq_m^3}{dt} = v_m^2 - v_m^3$$
$$\frac{dq_m^4}{dt} = v_m^3 - v_m^4 \qquad \frac{dq_m^5}{dt} = v_m^4 - v_m^5 \qquad \frac{dq_m^6}{dt} = v_m^5 - v_m^6$$
$$\frac{dq_m^7}{dt} = v_m^6 - v_m^7 \qquad \frac{dq_m^8}{dt} = v_m^7 - v_m^8 \qquad \frac{dq_m^9}{dt} = v_m^8 - v_m^9$$
$$\frac{dq_m^{10}}{dt} = v_m^9 - v_m^{10} \qquad \frac{dq_m^{11}}{dt} = v_m^{10} - v_m^{11} \qquad \frac{dq_m^{12}}{dt} = v_m^{11} - v_m^{12}$$
$$\frac{dq_m^{13}}{dt} = v_m^{12} - v_m^{13} \qquad \frac{dq_m^{14}}{dt} = v_m^{13} - v_m^{14} \qquad \frac{dq_m^{15}}{dt} = v_m^{14} - v_m^{15}$$

The chemical potentials are given by the following.

$$u_m^1 = RT\ln(K_1 q_m^1) \quad u_m^2 = RT\ln(K_2 q_m^2) \quad u_m^3 = RT\ln(K_3 q_m^3) \quad u_m^4 = RT\ln(K_4 q_m^4) \quad u_m^5 = RT\ln(K_5 q_m^5)$$
$$u_m^6 = RT\ln(K_6 q_m^6) \quad u_m^7 = RT\ln(K_7 q_m^7) \quad u_m^8 = RT\ln(K_8 q_m^8) \quad u_m^9 = RT\ln(K_9 q_m^9) \quad u_m^{10} = RT\ln(K_{10} q_m^{10})$$
$$u_m^{11} = RT\ln(K_{11} q_m^{11}) \quad u_m^{12} = RT\ln(K_{12} q_m^{12}) \quad u_m^{13} = RT\ln(K_{13} q_m^{13}) \quad u_m^{14} = RT\ln(K_{14} q_m^{14}) \quad u_m^{15} = RT\ln(K_{15} q_m^{15})$$

$$u_i^{K+} = RT\ln(K_i^K q_i^{K+}) \quad u_i^{Na+} = RT\ln(K_i^{Na} q_i^{Na+}) \quad u_o^{Na+} = RT\ln(K_o^{Na} q_o^{Na+}) \quad u_o^{K+} = RT\ln(K_o^K q_o^{K+})$$
$$u_i^{ATP} = RT\ln(K_i^{ATP} q_i^{ATP}) \quad u_i^{ADP} = RT\ln(K_i^{ADP} q_i^{ADP}) \quad u_i^{P_i} = RT\ln(K_i^{P_i} q_i^{P_i}) \quad u_i^H = RT\ln(K_i^H q_i^H)$$

The flow rates are given by:

$$v_m^1 = \kappa_1 \left( K_1 q_m^1 - K_2 q_m^2 K_i^K q_i^{K+} \right) \qquad v_m^2 = \kappa_2 \left( K_2 q_m^2 - K_3 q_m^3 K_i^K q_i^{K+} \right)$$

$$v_m^3 = \kappa_3 \left( K_3 q_m^3 K_i^{Na} q_i^{Na+} - K_4 q_m^4 \right) \qquad v_m^4 = \kappa_4 \left( K_4 q_m^4 K_i^{Na} q_i^{Na+} - K_5 q_m^5 \right)$$

$$v_m^5 = \kappa_5 \left( K_5 q_m^5 K_i^{Na} q_i^{Na+} - K_6 q_m^6 \exp\left( \frac{z_1 F u_m^e}{RT} \right) \right) \qquad v_m^6 = \kappa_6 \left( K_6 q_m^6 - K_7 q_m^7 K_i^{ADP} q_i^{ADP} \right)$$

$$v_m^7 = \kappa_7 \left( K_7 q_m^7 - K_8 q_m^8 \right) \qquad v_m^8 = \kappa_8 \left( K_8 q_m^8 - K_9 q_m^9 K_o^{Na} q_o^{Na+} \exp\left( \frac{z_2 F u_m^e}{RT} \right) \right)$$

$$v_m^9 = \kappa_9 \left( K_9 q_m^9 - K_{10} q_m^{10} K_o^{Na} q_o^{Na+} \right) \qquad v_m^{10} = \kappa_{10} \left( K_{10} q_m^{10} - K_{11} q_m^{11} K_o^{Na} q_o^{Na+} \right)$$

$$v_m^{11} = \kappa_{11} \left( K_{11} q_m^{11} K_o^K q_o^{K+} - K_{12} q_m^{12} \right) \qquad v_m^{12} = \kappa_{12} \left( K_{12} q_m^{12} K_o^K q_o^{K+} - K_{13} q_m^{13} \right)$$

$$v_m^{13} = \kappa_{13} \left( K_{13} q_m^{13} - K_{14} q_m^{14} K_i^{P_i} q_i^{P_i} K_i^H q_i^H \right) \qquad v_m^{14} = \kappa_{14} \left( K_{14} q_m^{14} K_i^{ATP} q_i^{ATP} - K_{15} q_m^{15} \right)$$

$$v_m^{15} = \kappa_{15} \left( K_{15} q_m^{15} - K_{16} q_m^{16} \right)$$

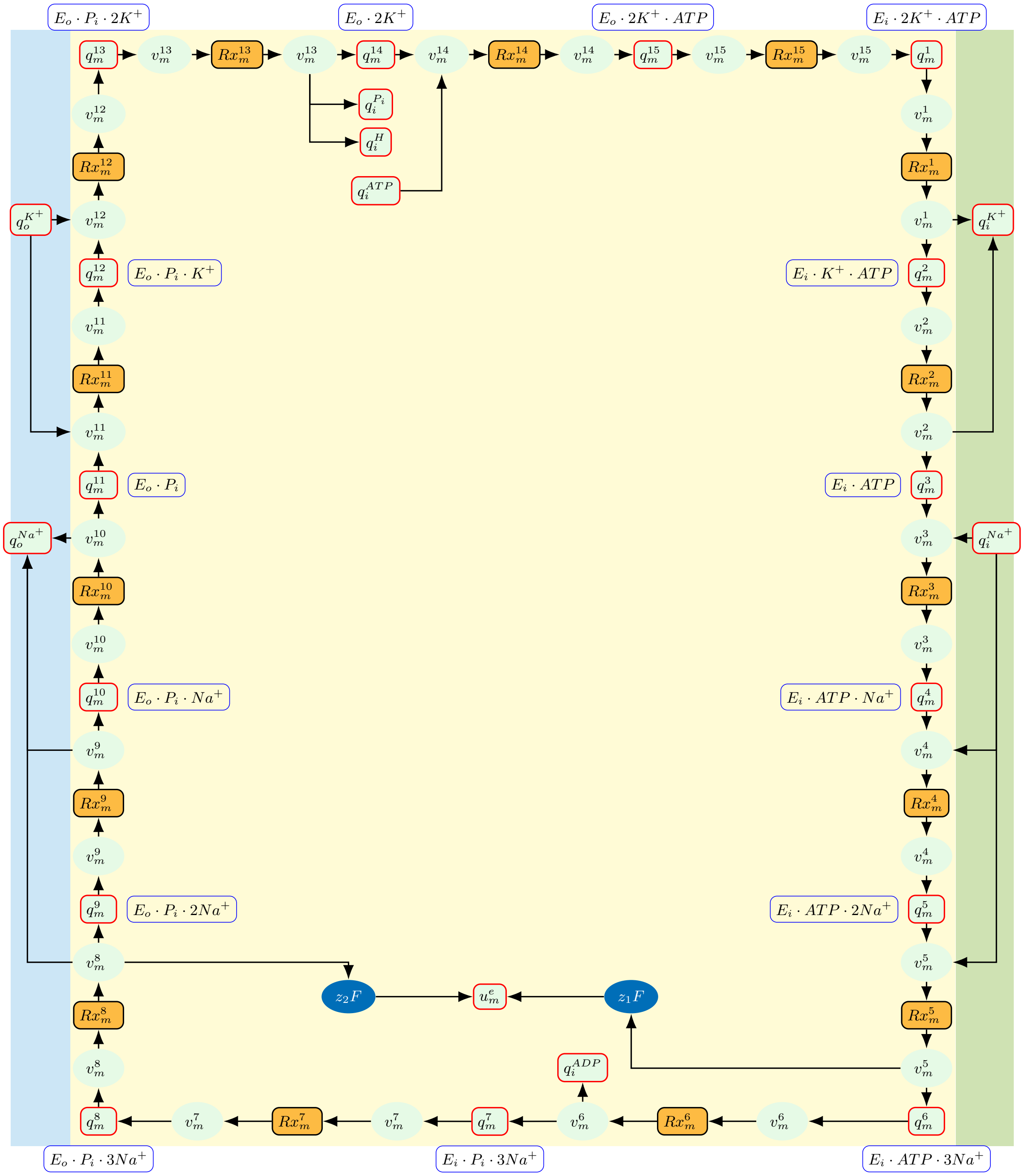


Figure 1: Bond graph of the 15-state model, adapted from (1).

Figure 2: Bond graph mode of NKA with 6 states.

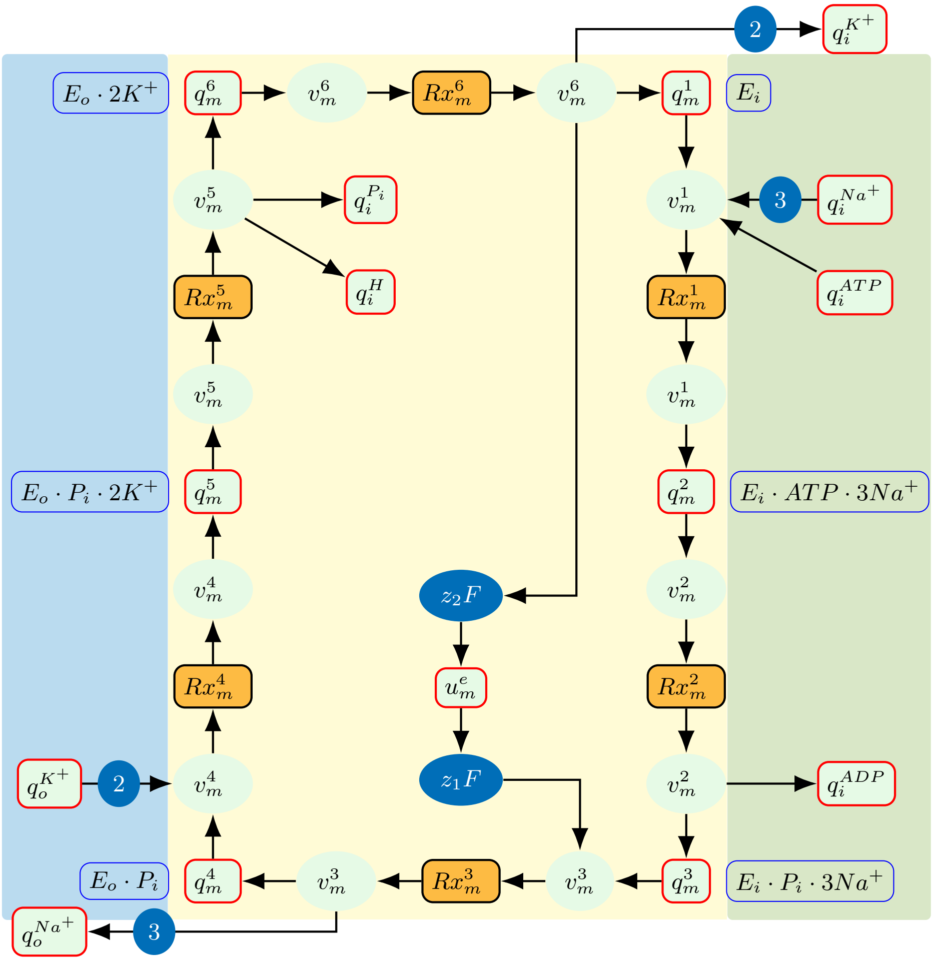


The CellML code of the model is *NKE_BG_15_state.cellml* and the parameters are in *NKE_BG_params.cellml*, while the file *NKE_BG_Env.cellml* contains the environmental variables and the membrane potentials used in the simulation. The simulation conditions are in the sedml files listed in *NKE_BG_15_state_sedmls.json*, while the python scripts to edit and run the sedml files are: *edit_sedmls_NKE_BG_15_state.py* and *run_sedmls_NKE_BG_15_state.py*.

The steady state flux of the 15-state model is given in the CellML code *Terkildsen_NaK_kinetic_modular.cellml* associated with the paper (1). We modified the code to allow the model access to the environmental variables in *NKE_BG_Env.cellml* and renamed it as *Terkildsen_NaK_kinetic_modular_V.cellml*. The simulation conditions are in the sedml files listed in *Terkildsen_NaK_kinetic_modular_V_sedmls.json*, while the python scripts to edit and run the sedml files are: *edit_sedmls_Terkildsen_NaK_kinetic.py* and *run_sedmls_Terkildsen_NaK_kinetic_modular_V.py*.

#### Bond graph modes of NKA with 6 states

The bond graph of the 6-state model is shown in Figure 2.

The equations for conservation of mass are expressed as below.

$$\frac{dq_m^1}{dt} = v_m^6 - v_m^1 \qquad \frac{dq_m^2}{dt} = v_m^1 - v_m^2 \qquad \frac{dq_m^3}{dt} = v_m^2 - v_m^3$$

$$\frac{dq_m^4}{dt} = v_m^3 - v_m^4 \qquad \frac{dq_m^5}{dt} = v_m^4 - v_m^5 \qquad \frac{dq_m^6}{dt} = v_m^5 - v_m^6$$

The chemical potentials are given by the following.

$$u_m^1 = RT\ln(K_1 q_m^1) \qquad u_m^2 = RT\ln(K_2 q_m^2) \qquad u_m^3 = RT\ln(K_3 q_m^3)$$
$$u_m^4 = RT\ln(K_4 q_m^4) \qquad u_m^5 = RT\ln(K_5 q_m^5) \qquad u_m^6 = RT\ln(K_6 q_m^6)$$

$$u_i^{K+} = RT\ln(K_i^K q_i^{K+}) \qquad u_i^{Na+} = RT\ln(K_i^{Na} q_i^{Na+}) \qquad u_o^{Na+} = RT\ln(K_o^{Na} q_o^{Na+}) \qquad u_o^{K+} = RT\ln(K_o^K q_o^{K+})$$
$$u_i^{ATP} = RT\ln(K_i^{ATP} q_i^{ATP}) \qquad u_i^{ADP} = RT\ln(K_i^{ADP} q_i^{ADP}) \qquad u_i^{P_i} = RT\ln(K_i^{P_i} q_i^{P_i}) \qquad u_i^{H} = RT\ln(K_i^{H} q_i^{H})$$

The flow rates are given by:

$$v_m^1 = \kappa_1\left(K_1 q_m^1 (K_i^{Na} q_i^{Na+})^3 K_i^{ATP} q_i^{ATP} - K_2 q_m^2\right) \qquad v_m^2 = \kappa_2\left(K_2 q_m^2 - K_3 q_m^3 K_i^{ADP} q_i^{ADP}\right)$$

$$v_m^3 = \kappa_3\left(K_3 q_m^3 \exp\left(\frac{z_1 F u_m^e}{RT}\right) - K_4 q_m^4 (K_o^{Na} q_o^{Na+})^3\right) \qquad v_m^4 = \kappa_4\left(K_4 q_m^4 (K_o^K q_o^{K+})^2 - K_5 q_m^5\right)$$

$$v_m^5 = \kappa_5\left(K_5 q_m^5 - K_6 q_m^6 K_i^{P_i} q_i^{P_i} K_i^H q_i^H\right) \qquad v_m^6 = \kappa_6\left(K_6 q_m^6 - K_7 q_m^7 (K_i^K q_i^{K+})^2 \exp\left(\frac{z_2 F u_m^e}{RT}\right)\right)$$

To derive an analytic formula for the steady state behaviour of the 6-state model, we make the assumption that the substrates are present in much higher quantities than the membrane-bound transporter (i.e., the Briggs-Haldane assumption (2)), and that the cycle is therefore transitioning through the six states at a constant steady-state rate $v$, i.e.,

$$v_m^1 = v_m^2 = v_m^3 = v_m^4 = v_m^5 = v_m^6 = v. \tag{1}$$

These equations are supplemented with the constraint on the total amount of protein:

$$q_{\text{tot}} = q_m^1 + q_m^2 + q_m^3 + q_m^4 + q_m^5 + q_m^6 \tag{2}$$

or

$$\frac{\bar{q}_m^1}{K_1} + \frac{\bar{q}_m^2}{K_2} + \frac{\bar{q}_m^3}{K_3} + \frac{\bar{q}_m^4}{K_4} + \frac{\bar{q}_m^5}{K_5} + \frac{\bar{q}_m^6}{K_6} = q_{\text{tot}} \tag{3}$$

where $\bar{q}_m^i = K_i q_m^i$ for $i = 1, 2, ..., 6$.

Rewriting the flow equations under the steady state assumption (Eq. 1) allows us to express the quantities of each state in terms of the steady state flux $v$.

$$\bar{q}_m^2 = \bar{q}_m^1 \cdot \left(\bar{q}_i^{\text{Na}^+}\right)^3 \cdot \bar{q}_i^{\text{ATP}} - \frac{v}{\kappa_1} \tag{4}$$

$$\bar{q}_m^3 = \frac{\bar{q}_m^2 - \frac{v}{\kappa_2}}{\bar{q}_i^{\text{ADP}}} \tag{5}$$

With Eq.4, we can express Eq.5 as:

$$\bar{q}_m^3 = \bar{q}_m^1 \cdot \left(\bar{q}_i^{\text{Na}^+}\right)^3 \cdot \frac{\bar{q}_i^{\text{ATP}}}{\bar{q}_i^{\text{ADP}}} - \frac{\frac{v}{\kappa_1} + \frac{v}{\kappa_2}}{\bar{q}_i^{\text{ADP}}} \tag{6}$$

$$\bar{q}_m^4 = \left(\bar{q}_m^3 e^{\frac{z_1 F u_m^e}{RT}} - \frac{v}{\kappa_3}\right)\left(\bar{q}_o^{\text{Na}^+}\right)^{-3} \tag{7}$$

With Eq.6, we can express Eq. 7 as:

$$\bar{q}_m^4 = \bar{q}_m^1 \left(\frac{\bar{q}_i^{\text{Na}^+}}{\bar{q}_o^{\text{Na}^+}}\right)^3 \frac{\bar{q}_i^{\text{ATP}}}{\bar{q}_i^{\text{ADP}}} e^{\frac{z_1 F u_m^e}{RT}} - \frac{\frac{v}{\kappa_1} + \frac{v}{\kappa_2}}{\bar{q}_i^{\text{ADP}}\left(\bar{q}_o^{\text{Na}^+}\right)^3} e^{\frac{z_1 F u_m^e}{RT}} - \frac{v}{\kappa_3\left(\bar{q}_o^{\text{Na}^+}\right)^3} \tag{8}$$

$$\bar{q}_m^5 = \bar{q}_m^4 \left(\bar{q}_o^{K^+}\right)^2 - \frac{v}{\kappa_4} \tag{9}$$

With Eq.8, we can express Eq.9 as:

$$\bar{q}_m^5 = \left[\bar{q}_m^1 \left(\frac{\bar{q}_i^{\mathrm{Na}^+}}{\bar{q}_o^{\mathrm{Na}^+}}\right)^3 \frac{\bar{q}_i^{\mathrm{ATP}}}{\bar{q}_i^{\mathrm{ADP}}} e^{\frac{z_1 F u_m^e}{RT}} - \frac{\frac{v}{\kappa_1} + \frac{v}{\kappa_2}}{\bar{q}_i^{\mathrm{ADP}} \left(\bar{q}_o^{\mathrm{Na}^+}\right)^3} e^{\frac{z_1 F u_m^e}{RT}} - \frac{v}{\kappa_3 \left(\bar{q}_o^{\mathrm{Na}^+}\right)^3}\right] \left(\bar{q}_o^{K^+}\right)^2 - \frac{v}{\kappa_4} \tag{10}$$

$$\bar{q}_m^6 = \frac{\bar{q}_m^5 - \frac{v}{\kappa_5}}{\bar{q}_i^{P_i} \cdot \bar{q}_i^{\mathrm{H}^+}} \tag{11}$$

With Eq.10, we can express Eq.11 as:

$$\bar{q}_m^6 = \left[\bar{q}_m^1 \left(\frac{\bar{q}_i^{\mathrm{Na}^+}}{\bar{q}_o^{\mathrm{Na}^+}}\right)^3 \frac{\bar{q}_i^{\mathrm{ATP}}}{\bar{q}_i^{\mathrm{ADP}} \cdot \bar{q}_i^{P_i} \cdot \bar{q}_i^{\mathrm{H}^+}} e^{\frac{z_1 F u_m^e}{RT}} - \frac{\frac{v}{\kappa_1} + \frac{v}{\kappa_2}}{\bar{q}_i^{\mathrm{ADP}} \cdot \bar{q}_i^{P_i} \cdot \bar{q}_i^{\mathrm{H}^+} \cdot \left(\bar{q}_o^{\mathrm{Na}^+}\right)^3} e^{\frac{z_1 F u_m^e}{RT}} - \frac{v}{\kappa_3 \cdot \bar{q}_i^{P_i} \cdot \bar{q}_i^{\mathrm{H}^+} \cdot \left(\bar{q}_o^{\mathrm{Na}^+}\right)^3}\right] \left(\bar{q}_o^{K^+}\right)^2 - \frac{\frac{v}{\kappa_4} + \frac{v}{\kappa_5}}{\bar{q}_i^{P_i} \cdot \bar{q}_i^{\mathrm{H}^+}} \tag{12}$$

From the expression of $v_m^6$, we get the relation:

$$\bar{q}_m^1 \, (\bar{q}_i^{K^+})^2 \, e^{\frac{z_2 F u_m^e}{RT}} = \bar{q}_m^6 - \frac{v}{\kappa_6} \tag{13}$$

Substituting $\bar{q}_m^6$ with Eq.12 in Eq.13, we get:

$$\bar{q}_m^1 \, (\bar{q}_i^{K^+})^2 \, e^{\frac{z_2 F u_m^e}{RT}} = \left[\bar{q}_m^1 \left(\frac{\bar{q}_i^{\mathrm{Na}^+}}{\bar{q}_o^{\mathrm{Na}^+}}\right)^3 \frac{\bar{q}_i^{\mathrm{ATP}}}{\bar{q}_i^{\mathrm{ADP}} \cdot \bar{q}_i^{P_i} \cdot \bar{q}_i^{\mathrm{H}^+}} e^{\frac{z_1 F u_m^e}{RT}} - \frac{\frac{v}{\kappa_1} + \frac{v}{\kappa_2}}{\bar{q}_i^{\mathrm{ADP}} \cdot \bar{q}_i^{P_i} \cdot \bar{q}_i^{\mathrm{H}^+} \cdot \left(\bar{q}_o^{\mathrm{Na}^+}\right)^3} e^{\frac{z_1 F u_m^e}{RT}} - \frac{v}{\kappa_3 \cdot \bar{q}_i^{P_i} \cdot \bar{q}_i^{\mathrm{H}^+} \cdot \left(\bar{q}_o^{\mathrm{Na}^+}\right)^3}\right] \left(\bar{q}_o^{K^+}\right)^2 - \frac{\frac{v}{\kappa_4} + \frac{v}{\kappa_5}}{\bar{q}_i^{P_i} \cdot \bar{q}_i^{\mathrm{H}^+}} - \frac{v}{\kappa_6} \tag{14}$$

Reorganizing Eq.14 gives:

$$\bar{q}_m^1 \left\{ \left(\frac{\bar{q}_i^{\mathrm{Na}^+}}{\bar{q}_o^{\mathrm{Na}^+}}\right)^3 \frac{\bar{q}_i^{\mathrm{ATP}}}{\bar{q}_i^{\mathrm{ADP}} \bar{q}_i^{\mathrm{P}_i} \bar{q}_i^{\mathrm{H}^+}} e^{\frac{z_1 F u_m^e}{RT}} \left(\bar{q}_o^{\mathrm{K}^+}\right)^2 - \left(\bar{q}_i^{\mathrm{K}^+}\right)^2 e^{\frac{z_2 F u_m^e}{RT}} \right\} = \left[ \frac{\frac{v}{\kappa_1} + \frac{v}{\kappa_2}}{\bar{q}_i^{\mathrm{ADP}} \bar{q}_i^{\mathrm{P}_i} \bar{q}_i^{\mathrm{H}^+} \left(\bar{q}_o^{\mathrm{Na}^+}\right)^3} e^{\frac{z_1 F u_m^e}{RT}} + \frac{\frac{v}{\kappa_3}}{\bar{q}_i^{\mathrm{P}_i} \bar{q}_i^{\mathrm{H}^+} \left(\bar{q}_o^{\mathrm{Na}^+}\right)^3} \right] \left(\bar{q}_o^{\mathrm{K}^+}\right)^2 + \frac{\frac{v}{\kappa_4} + \frac{v}{\kappa_5}}{\bar{q}_i^{\mathrm{P}_i} \bar{q}_i^{\mathrm{H}^+}} + \frac{v}{\kappa_6} \tag{15}$$

Multiplying trought by $e^{\frac{-z_1 F u_m^e}{RT}} \left(\bar{q}_o^{\mathrm{K}^+}\right)^{-2}$ and using the net charge movement relation $z_1 - z_2 = 1$, we can express the Eq. 15 as:

$$\bar{q}_m^1 \left\{ \left(\frac{\bar{q}_i^{\mathrm{Na}^+}}{\bar{q}_o^{\mathrm{Na}^+}}\right)^3 \frac{\bar{q}_i^{\mathrm{ATP}}}{\bar{q}_i^{\mathrm{ADP}} \bar{q}_i^{\mathrm{P}_i} \bar{q}_i^{\mathrm{H}^+}} - \left(\frac{\bar{q}_i^{\mathrm{K}^+}}{\bar{q}_o^{\mathrm{K}^+}}\right)^2 e^{\frac{-F u_m^e}{RT}} \right\} = \frac{\frac{v}{\kappa_1} + \frac{v}{\kappa_2}}{\bar{q}_i^{\mathrm{ADP}} \bar{q}_i^{\mathrm{P}_i} \bar{q}_i^{\mathrm{H}^+} \left(\bar{q}_o^{\mathrm{Na}^+}\right)^3} + \frac{\frac{v}{\kappa_3}}{\bar{q}_i^{\mathrm{P}_i} \bar{q}_i^{\mathrm{H}^+} \left(\bar{q}_o^{\mathrm{Na}^+}\right)^3} e^{\frac{-z_1 F u_m^e}{RT}} + \left[ \frac{\frac{v}{\kappa_4} + \frac{v}{\kappa_5}}{\bar{q}_i^{\mathrm{P}_i} \bar{q}_i^{\mathrm{H}^+}} + \frac{v}{\kappa_6} \right] \left(\bar{q}_o^{\mathrm{K}^+}\right)^{-2} e^{\frac{-z_1 F u_m^e}{RT}} \tag{16}$$

Rearranging the terms gives an expression for $\bar{q}_m^1$:

$$\bar{q}_m^1 = \frac{1}{A} \left\{ \frac{\frac{v}{\kappa_1} + \frac{v}{\kappa_2}}{\bar{q}_i^{\mathrm{ADP}} \bar{q}_i^{\mathrm{P}_i} \bar{q}_i^{\mathrm{H}^+} \left(\bar{q}_o^{\mathrm{Na}^+}\right)^3} + \frac{\frac{v}{\kappa_3}}{\bar{q}_i^{\mathrm{P}_i} \bar{q}_i^{\mathrm{H}^+} \left(\bar{q}_o^{\mathrm{Na}^+}\right)^3} e^{\frac{-z_1 F u_m^e}{RT}} + \left[ \frac{\frac{v}{\kappa_4} + \frac{v}{\kappa_5}}{\bar{q}_i^{\mathrm{P}_i} \bar{q}_i^{\mathrm{H}^+}} + \frac{v}{\kappa_6} \right] \left(\bar{q}_o^{\mathrm{K}^+}\right)^{-2} e^{\frac{-z_1 F u_m^e}{RT}} \right\} \tag{17}$$

where

$$A = \left(\frac{\bar{q}_i^{\mathrm{Na}^+}}{\bar{q}_o^{\mathrm{Na}^+}}\right)^3 \frac{\bar{q}_i^{\mathrm{ATP}}}{\bar{q}_i^{\mathrm{ADP}} \bar{q}_i^{\mathrm{P}_i} \bar{q}_i^{\mathrm{H}^+}} - \left(\frac{\bar{q}_i^{\mathrm{K}^+}}{\bar{q}_o^{\mathrm{K}^+}}\right)^2 e^{\frac{-F u_m^e}{RT}} \tag{18}$$

Substituting $\bar{q}_m^1$ (Eq.17), $\bar{q}_m^2$ (Eq.4), $\bar{q}_m^3$ (Eq.6), $\bar{q}_m^4$ (Eq.8), $\bar{q}_m^5$ (Eq.10), and $\bar{q}_m^6$ (Eq.12) into the total protein constraint (Eq.3) allows us to solve for the steady state flux $v$. To simplify the analysis, we define the following:

$$\Lambda_i = \frac{K_6}{K_i}, \quad \text{for } i = 1, 2, 3, 4, 5 \tag{19}$$

After rearranging the terms and simplifying, the final expression for the steady state flux of the 6-state model is given by:

$$v^{\mathrm{NKE}} = \frac{K_6 \kappa_6 q_{\mathrm{tot}}}{C - AD} A \tag{20}$$

where

$$A = \left(\frac{\bar{q}_i^{\mathrm{Na^+}}}{\bar{q}_o^{\mathrm{Na^+}}}\right)^3 \frac{\bar{q}_i^{\mathrm{ATP}}}{\bar{q}_i^{\mathrm{ADP}}\,\bar{q}_i^{P_i}\,\bar{q}_i^{H}} - \left(\frac{\bar{q}_i^{\mathrm{K^+}}}{\bar{q}_o^{\mathrm{K^+}}}\right)^2 e^{-\frac{Fu_m^e}{RT}} \tag{21}$$

$$B = \left(\frac{\bar{q}_i^{\mathrm{Na^+}}}{\bar{q}_o^{\mathrm{Na^+}}}\right)^3 \frac{\bar{q}_i^{\mathrm{ATP}}}{\bar{q}_i^{\mathrm{ADP}}\,\bar{q}_i^{P_i}\,\bar{q}_i^{H}}\, e^{\frac{z_1 Fu_m^e}{RT}} \left[\Lambda_4 + \left(\Lambda_5 + \frac{1}{\bar{q}_i^{P_i}\,\bar{q}_i^{H}}\right)\left(\bar{q}_o^{\mathrm{K^+}}\right)^2\right] \tag{22}$$

$$\begin{aligned} C = {} & \left[\frac{\Lambda_1}{\bar{q}_i^{P_i}\,\bar{q}_i^{H^+}} + \Lambda_2 \frac{\left(\bar{q}_i^{\mathrm{Na^+}}\right)^3 \bar{q}_i^{\mathrm{ATP}}}{\bar{q}_i^{P_i}\,\bar{q}_i^{H^+}} + \Lambda_3 \frac{\left(\bar{q}_i^{\mathrm{Na^+}}\right)^3 \bar{q}_i^{\mathrm{ATP}}}{\bar{q}_i^{\mathrm{ADP}}\,\bar{q}_i^{P_i}\,\bar{q}_i^{H^+}} + B\right] \\ & \times \left[\frac{\lambda_1 + \lambda_2}{\bar{q}_i^{\mathrm{ADP}}\left(\bar{q}_o^{\mathrm{Na^+}}\right)^3} + \frac{\lambda_3}{\left(\bar{q}_o^{\mathrm{Na^+}}\right)^3} \exp\left(-\frac{z_1 Fu_m^e}{RT}\right) \right. \\ & \left. + \left(\lambda_4 + \lambda_5 + \bar{q}_i^{P_i}\,\bar{q}_i^{H^+}\right)\left(\bar{q}_o^{\mathrm{K^+}}\right)^{-2} \exp\left(-\frac{z_1 Fu_m^e}{RT}\right)\right]. \end{aligned} \tag{23}$$

$$\begin{aligned} D = {} & \Lambda_2\lambda_1 + \Lambda_5\lambda_4 + \frac{\lambda_4 + \lambda_5}{\bar{q}_i^{P_i}\,\bar{q}_i^{H^+}} + \Lambda_3 \frac{\lambda_1 + \lambda_2}{\bar{q}_i^{\mathrm{ADP}}} \\ & + \left(\frac{\lambda_1 + \lambda_2}{\bar{q}_i^{\mathrm{ADP}}} \exp\left(\frac{z_1 Fu_m^e}{RT}\right) + \lambda_3\right) \frac{1}{\left(\bar{q}_o^{\mathrm{Na^+}}\right)^3} \left[\Lambda_4 + \left(\Lambda_5 + \frac{1}{\bar{q}_i^{P_i}\,\bar{q}_i^{H^+}}\right)\left(\bar{q}_o^{\mathrm{K^+}}\right)^2\right]. \end{aligned} \tag{24}$$

$\lambda_i = \frac{\kappa_6}{\kappa_i}$, and $\Lambda_i = \frac{K_6}{K_i}$ for $i = 1, 2, ..., 5$.

$$\bar{q}_i^{\mathrm{K^+}} = K_i^K q_i^{K+} \qquad \bar{q}_i^{\mathrm{Na^+}} = K_i^{Na} q_i^{Na+} \qquad \bar{q}_o^{\mathrm{Na^+}} = K_o^{Na} q_o^{Na+} \qquad \bar{q}_o^{\mathrm{K^+}} = K_o^K q_o^{K+}$$

$$\bar{q}_i^{\mathrm{ATP}} = K_i^{ATP} q_i^{ATP} \qquad \bar{q}_i^{\mathrm{ADP}} = K_i^{ADP} q_i^{ADP} \qquad \bar{q}_i^{P_i} = K_i^{P_i} q_i^{P_i} \qquad \bar{q}_i^{H} = K_i^H q_i^H$$

Table 1 lists the parameters used in the 6-state model.
Table 2 lists and describes the files related to the 6-state model.

#### Instructions to run the simulations and reproduce the figures

The CellML and sedml files are in workspace `https://models.physiomeproject.org/workspace/cad`, while the python scripts are in Github repository `https://github.com/WeiweiAi/EA_NKE`. Since the Github repository includes sumbodules, please clone the repository using the command:

```
git clone https://github.com/WeiweiAi/EA_NKE.git --recurse-submodules
```

It is recommended to use Python version 3.11. Please create a virtual environment and install the required packages using the command:

```
pip install -r requirements.txt
```

Table 1: Parameters used in the 6-state model.

| Parameter | Value | Unit |
|---|---|---|
| $\kappa_1$ | 62051.4 | $fmol \cdot s^{-1}$ |
| $\kappa_2$ | 0.549906 | $fmol \cdot s^{-1}$ |
| $\kappa_3$ | 474.926 | $fmol \cdot s^{-1}$ |
| $\kappa_4$ | 910019 | $fmol \cdot s^{-1}$ |
| $\kappa_5$ | 384.851 | $fmol \cdot s^{-1}$ |
| $\kappa_6$ | 3412.49 | $fmol \cdot s^{-1}$ |
| $K_1$ | 3.64797 | $fmol^{-1}$ |
| $K_2$ | 80.1178 | $fmol^{-1}$ |
| $K_3$ | 99322.9 | $fmol^{-1}$ |
| $K_4$ | 49.0681 | $fmol^{-1}$ |
| $K_5$ | 84.2138 | $fmol^{-1}$ |
| $K_6$ | 727440 | $fmol^{-1}$ |
| $z_1$ | 0.945049 | dimensionless |
| $z_2$ | -0.054951 | dimensionless |

Table 2: Description of the files related to the 6-state model.

| File name | Description |
|---|---|
| *NKE_BG_6_state_ATPNaZKV2.cellml* | CellML code of the 6-state model |
| *NKE_BG_6_state_ATPNaZKV2_SS.cellml* | CellML code of the steady state flux of the 6-state model |
| *NKE_BG_6_state_ATPNaZKV2_fixedV.cellml* | CellML code of the 6-state model with fixed membrane potential |
| *NKE_BG_6_state_ATPNaZKV2_SS_fixedV.cellml* | CellML code of the steady state flux of the 6-state model with fixed membrane potential |
| *NKE_BG_6_state_ATPNaZKV2_param.cellml* | Parameters for the 6-state model |
| *NKE_BG_6_state_ATPNaZKV2_sedmls.json* | The list of sedml files for the 6-state model |
| *NKE_BG_6_state_ATPNaZKV2_fixedV_sedmls.json* | The list of sedml files for the 6-state model with fixed membrane potential |
| *NKE_BG_6_state_ATPNaZKV2_SS_sedmls.json* | The list of sedml files for the steady state flux of the 6-state model |
| *NKE_BG_6_state_ATPNaZKV2_SS_fixedV_sedmls.json* | The list of sedml files for the steady state flux of the 6-state model with fixed membrane potential |
| *edit_sedmls_NKE_BG_6_state_ATPNaZKV2.py* | Python script to edit the sedml files for the 6-state model |
| *run_sedmls_NKE_BG_6_state_ATPNaZKV2.py* | Python script to run the sedml files for the 6-state model |

Please move to the scripts directory: *cd scripts* and the procedures to run the simulations and reproduce the figures are described below:

1. *python run_sedmls_Terkildsen_kinetic.py* to get the data of the steady state flux of the 15-state model under varying conditions.

2. *python run_sedmls_NKE_BG_15_state.py* to get the data of the full bond graph of the 15-state model under varying conditions.

3. *python run_sedmls_NKE_BG_6_state_ATPNaZKV2.py* to run the simulations of the 6-state models under varying conditions.

4. *python EA_calc_6stateV2.py* to calculate the energetic quantities of the 6-state model under varying conditions.

5. *python EA_calc_15state.py* to calculate the energetic quantities of the 15-state model under varying conditions.

6. *python plot_original_ssV2.py* to reproduce Figure 4 in the main paper.

7. *python plot_EA_flow_6stateV2.py*, *python plot_EA_flow_15state.py* and *python plot_EA_bar_heatV2.py* to reproduce Figure 5 in the main paper.

8. *python plot_EA_2D_combine15state.py* and *python plot_EA_2D_combine6stateV2.py* to reproduce Figure 6 in the main paper.

9. *python plot_EA_2D_combine15state_deltaATP.py* and *python plot_EA_2D_combine6stateV2_deltaATP.py* to reproduce Figures 7,8 and 9 in the main paper.

10. *python plot_EA_2D_combine15state_flowRate.py* and *python plot_EA_2D_combine6stateV2_flowRate.py* to reproduce Figure 10 in the main paper.

11. *python plot_EA_2D_combine15state_NaK.py* and *python plot_EA_2D_combine6stateV2_NaK.py* to reproduce Figure 10 in the main paper.

#### Parameters used in the 6-state model for enterocyte simulation

Table 3: Parameters used in the 6-state model for enterocyte simulation.

| Parameter | Value | Unit |
|---|---|---|
| $\kappa_1$ | 2838.89 | $fmol \cdot s^{-1}$ |
| $\kappa_2$ | 51.10 | $fmol \cdot s^{-1}$ |
| $\kappa_3$ | 469.35 | $fmol \cdot s^{-1}$ |
| $\kappa_4$ | 619068.3 | $fmol \cdot s^{-1}$ |
| $\kappa_5$ | 35.87 | $fmol \cdot s^{-1}$ |
| $\kappa_6$ | 868.36 | $fmol \cdot s^{-1}$ |
| $K_1$ | 0.01673 | $fmol^{-1}$ |
| $K_2$ | 4477.02 | $fmol^{-1}$ |
| $K_3$ | 46639.87 | $fmol^{-1}$ |
| $K_4$ | 720.58 | $fmol^{-1}$ |
| $K_5$ | 1.6069 | $fmol^{-1}$ |
| $K_6$ | 30254.3 | $fmol^{-1}$ |
| $z_1$ | 0.945049 | dimensionless |
| $z_2$ | -0.054951 | dimensionless |
